# Intrinsic space-time couplings governing multi-scale cortical dynamics

**DOI:** 10.64898/2026.05.27.726038

**Authors:** Alexander D. White, Yu Wang, John Kochalka, Yukun Alex Hao, Chelsea Li, Melissa Hernandez, Nadya Andini, Ki Eun Pyo, Wendy Wenderski, Anish Mitra, Barsin Eshaghi Gharagoz, Chandan Kadur, Surya Ganguli, Thomas R. Clandinin, Michael Z. Lin, Karl Deisseroth

## Abstract

The neocortex covers a vast expanse of the mammalian brain and represents the principal target of clinical neuromodulation; however, the global principles of neocortical operation have been challenging to identify. In this regard, a limitation has been tracking activity with cortex-wide spatial coverage while maintaining access to the millisecond temporal resolution of neuronal firing— a crucial combination not achievable with existing recording technologies. Here we introduce and apply conformal immersion microscopy, enabling activity tracking across the entire dorsal cortex with millisecond temporal resolution and 100 µm spatial resolution (thus spanning five orders of magnitude in time and four in space), at sufficient sensitivity to resolve single-trial activity beyond 100 Hz. Drawing on physics-based frameworks, we apply multiscale analysis to identify a fundamental frequency-dependent coherence length that partitions the neocortex into discrete dynamical elements with well-defined propagation speeds, boundaries, and scale-invariant dynamics. These dynamical elements were found to be conserved from sub-threshold to suprathreshold (neuronal firing) regimes of neural activity, and were robust to diverse pharmacological, optogenetic, and genetic interventions. However, it was possible to identify and establish conditions allowing elemental boundaries to be selectively overridden, and to allow perturbation of specific elements even while conserving global dynamical architecture. Together, these findings enable measurement of intrinsic spatiotemporal parameters governing the dynamical organization of neocortex, which may provide a foundation for mechanistically-informed basic and translational understanding.

## Introduction

Elucidating the dynamical organization of the intact functioning neocortex (the largest, and most recently-and rapidly-evolving, structure in the mammalian brain) will likely be essential for understanding the neural underpinnings of complex adaptive behaviors and neuropsychiatric diseases. However, capturing cortex-spanning dynamics (with sufficient sensitivity and at relevant temporal and temporal scales) remains technically challenging, as current recording technologies are constrained by fundamental trade-offs among spatial coverage, temporal resolution, and signal-to-noise ratio (SNR). High-density electrophysiology provides the millisecond fidelity required to resolve local electrical fluctuations but lacks the coverage to capture continuous cortex-wide propagation of dynamics; in contrast, functional MRI (fMRI) offers global coverage but is limited by the slow, indirect nature of blood-oxygenation signals^1–5^, and electroencephalography (EEG) achieves whole-brain millisecond resolution but suffers from poor spatial resolution and coverage^6–9^. Wide-field optical encephalography (OEG)^10–12^ with genetically encoded voltage indicators (GEVIs)^13–17^ has emerged as a promising bridge into these electrical dynamics at the wide-field or mesoscopic scale^18–21^. Yet, traditional mesoscopic optics suffer from low light-collection efficiency, necessitating either long integration times that sacrifice temporal resolution or high-intensity illumination that leads to rapid photobleaching. Furthermore, the thin, flat focal planes of conventional systems fail to maintain focus across the curved geometry of the mammalian neocortex, which has prevented the capture of long-range (cortex-spanning) millisecond-scale activity maps.

Here we present and apply a technological and analytical framework – conformal immersion microscopy (CIM)– to maximize fast photon collection across large curved cortical areas, and we couple CIM to an advanced GEVI with high brightness and subthreshold sensitivity. By achieving high SNRs at low illumination intensities, this approach enables stable recording and control of electrical activity across the neocortex with unprecedented spatiotemporal resolution and duration, revealing a multi-scale functional architecture underlying dynamical operation of the intact mammalian neocortex.

### Ultrasensitive conformal imaging of cortical voltage dynamics

To enable fast activity imaging across cortex with high SNR and high spatial resolution, we developed conformal immersion microscopy (CIM) (summarized in **Fig. 1a-c, Extended Data Fig 1a-d**). This technique, building from tandem-lens widefield microscopy, surpasses the limits of conventional tandem-lens microscopy in two key ways. First, the low collection efficiency of tandem-lens microscopy was remedied with speed-boosters which trade off aperture for field of view (FOV)^22–24^, enabling a numerical aperture (NA) of 0.6 across a macroscopic 15mm FOV. However, this high NA exacerbates the second key issue with tandem-lens microscopy, namely that the thin flat focal field cannot capture the curved surface of cortex **(Fig 1b-c, Extended Data Fig 1d)**. We resolved this challenge by immersion-coupling the intact skull to a custom field-flattening lens designed to conform the focal plane to the cortical geometry. The lens curves the field with minimal optical aberrations, and the high-index glycerol immersion extends the effective depth of focus; the resulting system achieves uniform, high-resolution imaging across dorsal cortex, far exceeding the performance of uncorrected widefield optics **(Fig. 1c)**. Importantly, because CIM utilizes affordable components and standard immersion media, this method now provides an accessible platform that can be readily adapted to other existing mesoscopic systems.

**Figure 1:**
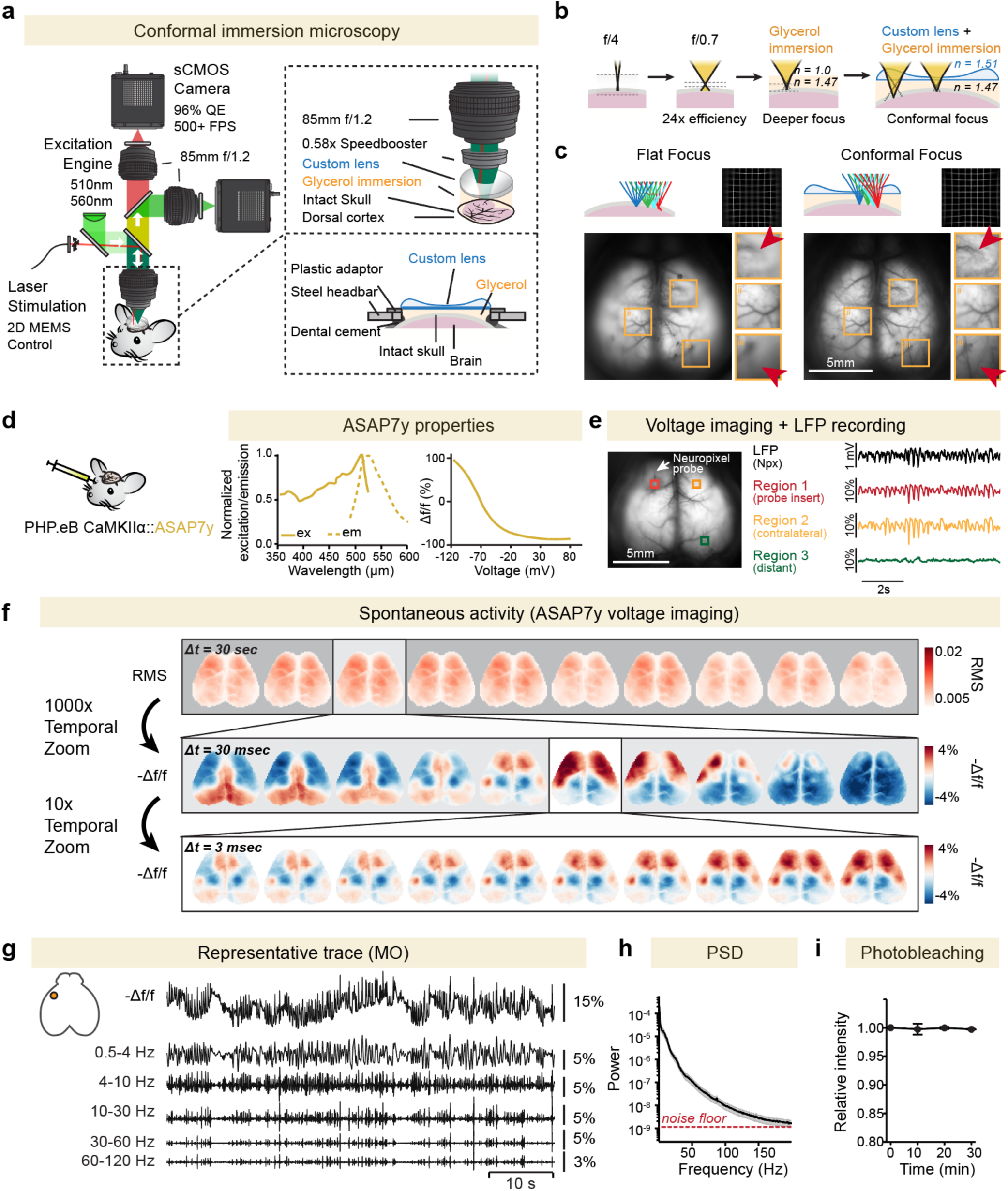
Sensitive and fast cortex-wide optical electrophysiology with conformal immersion microscopy. **a.** Overview of cortex-wide conformal immersion microscopy. Tandem lens microscopy enhanced by speed-booster lens, glycerol immersion, and custom plastic lens to conform the focal field to the cortical surface through the intact skull. **b.** Optical principles of conformal immersion. High numerical aperture (NA) increases photon collection but restricts depth of field. High-index glycerol immersion extends depth of focus, while a curved refractive interface aligns focal plane with cortical structure. **c.** Comparison of wide-field cortical imaging using a standard 0.6 NA objective (left) versus conformal immersion (right). Arrows indicate features resolved only after field-curvature correction. Scale bar, 5 mm. **d.** Viral expression strategy and voltage-sensor properties. Left: Pan-cortical ASAP7y expression via retro-orbital injection of AAV-PHP.eB CaMKIIα::ASAP7y. Right: Fluorescence spectra and voltage sensitivity (ΔF/F vs V_m_) of ASAP7y (adapted from Hao et al.^25^). **e.** Validation of optical signals via Neuropixels electrophysiological recording. Representative traces showing close correlation (r = 0.81 local, 0.76 contralateral, 0.18 distant site) between local ASAP7y fluorescence and simultaneous local field potential (LFP) recordings obtained via Neuropixels 1.0 probes (N = 2 mice). **f.** Multi-scale spontaneous voltage activity. High signal-to-noise ratio (SNR) from this approach enables single-trial imaging over five orders of magnitude in time without averaging (sampling rate, 333 Hz). **g.** Representative voltage trace decomposed into canonical frequency bands, demonstrating signal recovery up to the high-gamma range (60-120 Hz). Inset shows recording site in anterolateral cortex. **h.** Summary data quantifying power spectral density (PSD) of cortical voltage signals with this approach. Signal remains above the noise floor beyond 150Hz; noise floor is estimated by the frequency-independent floor of the PSD, reflecting shot-noise limit. Mean ± s.e.m.; N=4 mice, n=25 recording sessions. **i.** Negligible photobleaching observed over 30-min continuous recording. Mean ± s.e.m.; N=4 mice, n=25 recording sessions.

To leverage this enhanced light-collection capacity for high-speed voltage imaging, we expressed the novel bright inverse GEVI ASAP7y^25^ broadly across excitatory cortical neurons via retro-orbital injection **(Fig. 1d, Extended Data Fig. 1e-f)**. We first validated the fidelity of resulting optical signals via simultaneous Neuropixels recording (**Fig 1e**). Local ASAP7y fluorescence exhibited high correlation to local field potentials (LFP) (0.8 ± 0.014)^26^ with an order of magnitude higher coherence than that achieved by previously-reported GEVIs^15,21^ **(Fig. 1f, Extended Data 1n)**.This approach furthermore was found to enable the recording of spontaneous neural activity with unprecedented spatiotemporal resolution—130 µm pixels at 3ms sampling rate—providing sufficient SNR to enable resolution of clear single-trial dynamics **(Fig. 1f, Supplementary Video 1)** and measurement of cortical activity extending into the high gamma band (60–120 Hz) **(Fig. 1g)**, with neural activity events resolvable at frequencies exceeding 150 Hz **(Fig. 1h).**

The high collection efficiency of CIM, combined with next-generation indicator brightness, enables continuous imaging at high spatiotemporal resolution and high SNR with no discernable photobleaching over 30 min **(Fig. 1i)**. Collectively, these results establish the spatiotemporal precision and coverage necessary to resolve cortex-wide electrical dynamics previously inaccessible at this scale. CIM, here in combination with next-generation GEVIs, may provide new opportunities in the capturing of fast global voltage dynamics.

### Measuring dynamical parameters governing spontaneous cortical activity

We set out to explore spontaneous cortical activity during quiet wakefulness, leveraging the scope, speed, and sensitivity of this new approach. Spontaneous dynamics measured in this way across dorsal cortex revealed discrete, high-amplitude epochs spanning a broad frequency range (representative single-pixel spectrograms shown in **Extended Data Fig. 2a-e)**. When ranking time points by total spectral power, the majority of cortical power greater than ∼2–3 Hz was concentrated in <25% of the recording duration, while infralow activity (<2Hz) was more uniformly distributed in time **(Extended Data Fig. 2b)**. We began by mapping of interhemispheric coordination of these dynamics with CIM.

While homotopic regions (paired cortical regions in opposite hemispheres) were found to exhibit robust synchrony at low frequencies, these same homotopic pairs were found to desynchronize at high frequencies **(Fig. 2a, Extended Data Fig. 2f)**. Analysis of homotopic correlations in rolling 1-s windows revealed not only a component of zero-lag synchrony but also temporal structure **(Fig. 2b-c; Extended Data Fig. 2g-h)** marked by a bimodal distribution of consistent delay in which one hemisphere reliably led the other by ∼5-10 ms **(Extended Data Fig 2g-h)**. This delay is consistent with conduction speed of long-range myelinated callosal axons (∼2-5 mm/ms^27^) given that transcortical and interhemispheric axon path lengths in the mouse brain will reach several millimeters in length; the magnitude, consistency, and timescale of the 5-10 ms delay together suggest that interhemispheric coordination at these scales may be governed in large part by direct projections between cortical regions (in contrast to shared simultaneous drive from non-cortical structures which may contribute to the significant zero-lag component **(Fig. 2b)**).

**Figure 2:**
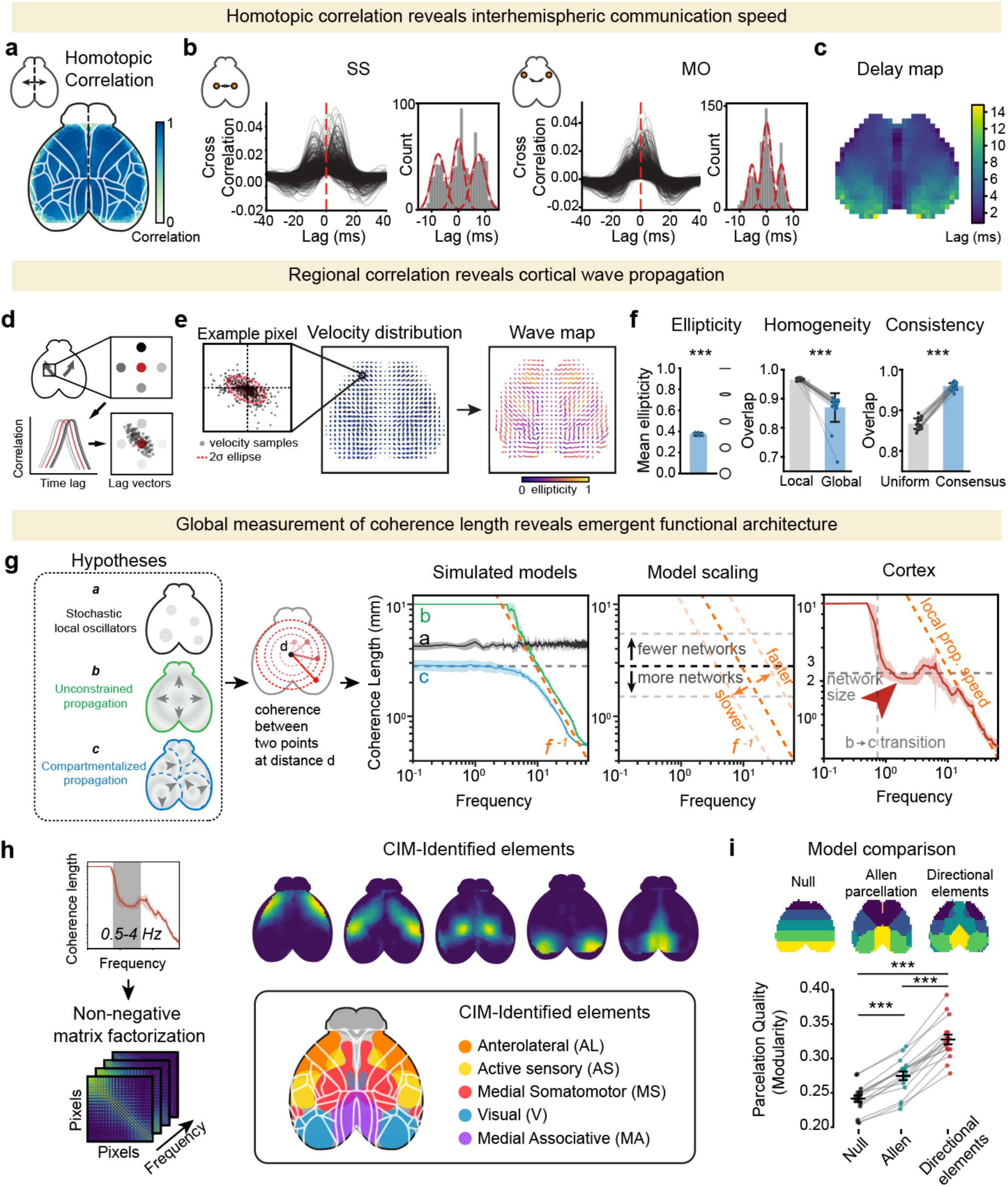
Emergent spatiotemporal global structure of spontaneous cortical dynamics. **a.** Resolving sub-20 ms latencies for inter-hemispheric communication (panels **a-c**). Panel **a** shows mean pixel-wise homotopic correlation (i.e. correlation between regions corresponding across hemispheres; N = 4 mice, n = 25 sessions); note that spontaneous activity between corresponding bilateral regions exhibits spatially homogeneous high correlation across dorsal cortex. **b.** Sub-second fluctuations in bilateral coordination. Rolling 1-s cross-correlation windows reveal non-simultaneous correlation peaks in spontaneous activity, with a symmetric delay distribution. Representative histograms highlight distinct peak timings between somatosensory (SS) and motor (MO) cortices; note the relatively greater zero-lag contribution in MO cortex. Histograms show combined peak times across N = 4 mice and n = 25 recording sessions; dashed lines show best fit to a symmetric trimodal normal distribution. Also see Extended Data Fig 2. **c.** Summary data for cortical mapping of mean homotopic delays, with measurement and mapping of intercortical timing enabled by the speed, sensitivity, and simultaneity of the imaging approach; heatmap shows mean homotopic delay maps across N= 4 mice in n = 25 recording sessions. Note that zero-lag contribution increases along the posterior-to-anterior axis. **d.** Vectorial analysis of spontaneous activity reveals structured cortical wave trajectories (panels **d-f**). Panel **d** shows local wave propagation analysis. To characterize local wave propagation (left), a lag vector was defined based on the direction of the peak cross-correlation lag in the neighborhood of a point. Cross-correlations were computed from 1-s rolling windows between each pixel and its four cardinal neighbors (top right, windows with minimal activity excluded). The resulting x–y cross-correlations (center right) defined a lag vector window. The distribution of these lag vectors (bottom right) reflects the directionality of wave propagation at this pixel. PCA of these distributions yielded ellipses that visualize the dominant propagation directions. **e.** Representative propagation bias and cortical wave maps. Velocity distribution for a representative single pixel in the motor cortex (left) illustrates the local propagation bias. The global map (middle) shows these velocity distributions across the cortex; vector length inversely represents wave propagation speed (i.e. longer vectors indicate slower wave propagation) while squeezed or elongated distributions reveal the dominant propagation mode along the principal axis. The wave map (right) further illustrates the primary propagation vectors with ellipticity (color scale) quantifying strength of directional dominance. **f.** Quantitative assessment of wave anisotropy and homogeneity (N = 4 mice, n =21 sessions). *Left*: Mean ellipticity is significantly greater than 0 (p < 0.001, one-tailed t-test), suggesting that locally there are dominant propagation directions. *Middle*: Local homogeneity (Bhattacharyya coefficient, see methods) of wave distributions is higher than global mean overlap (p < 0.001, paired t-test), showing that the direction of propagation is more locally consistent. *Right*: Whole-wave distribution consistency across mice sessions exceeds a homogeneous null model (p < 0.001, paired t-test), showing that the structure is conserved across animals (N = 4). Also see Extended Data Fig 3. **g.** Coherence length and the emergence of functional architecture (panels **g-j**). Panel **g** shows interaction of spatial and temporal scales of cortical activity. *Left*: Schematic of three dynamical model classes (stochastic local oscillators (i), unconstrained propagation (ii), and compartmentalized propagation (iii). *Middle*: Schematic of coherence length derivation. Coherence is calculated between pairs of points at increasing spatial separation (*d*), with coherence length defined as the maximum *d* maintaining a value above 1/e. *Right*: Coherence length versus frequency for null models, model scaling, and experimental data. Network count scales the vertical asymptote, and propagation velocity scales the 1/f asymptotic roll-off. Cortical data showed that the coherence length covers the entire cortex at low frequencies (<1Hz), decreases to about 2mm around 1Hz, and rolls off with a velocity of approximately 25mm/s above 12Hz. Data plotted as mean ± s.e.m. N = 4 mice, n = 25 sessions. **h.** Network-scale transition. Non-negative matrix factorization (NMF) of all-to-all pixel-wise coherence (1–10 Hz) reveals five elemental cortical networks. AL: anterolateral; AS: active sensing. MS: medial somatomotor; V: visual; MA: medial associative. Representative coherence matrix and NMF decomposition from one of N = 4 mice. **i.** Parcellation performance. NMF-derived networks provide significantly better data fits than both a naïve null model and traditional cytoarchitectural boundaries (Allen Mouse Brain Atlas; N = 4 mice, n = 16 sessions, paired t-test). Quality of network parcellation, measured by modularity^76^, is calculated for each session for each parcellation. Dynamic element parcellations were derived independently from the datasets used for evaluation.

We next explored mapping of intra-hemispheric activity patterns over space and time. Within each hemisphere, we derived local correlation lag vectors and quantified their spatial distribution via ellipticity, which measures the directional bias (anisotropy) of wave propagation. This analysis revealed that activity followed organized, bilaterally symmetrical spacetime trajectories **(Fig. 2d-e, Extended Data Fig. 3a-b)**. These trajectories were locally homogenous and significantly biased along principal directions, defining a stable dynamical topology that was highly conserved across both individuals and sessions **(Fig. 2f)**. The high degree of organization in both the direction and velocity of these waves revealed that spontaneous cortical activity across the scope of dorsal cortex followed orderly spatiotemporal dynamics^28–31^. Together with the propagation speed measurements in **Fig. 2a-c**, the measurement of these consistent trajectories suggested that the cortical surface, at the global scale, can be viewed as an organized medium governing the flow of fast electrical signaling.

To determine how temporal and spatial scales interact, we defined and measured cortical coherence length as the average distance over which two cortical points maintain coherence (> 1/e) **(Fig. 2g)**. To interpret the scaling of this length across frequencies, we charted the space of possibilities by evaluating three theoretical regimes: (a) a random-event model wherein coherence is frequency-independent, (b) a uniform medium wherein constant-velocity propagation results in brain-wide coherence at low frequencies followed by a 1/f roll-off, and (c) a partitioned network wherein anatomical boundaries restrict wave propagation, leading to a coherence length plateau corresponding to a characteristic scale of the network before transitioning to a 1/f roll-off at higher frequencies **(Fig. 2g, Extended Data Fig. 3c, Sup. Video 2)**. The plateau level reflects the number of distinct networks (fewer networks produce a higher plateau, more networks a lower one), while the slope of the high-frequency roll-off reflects local propagation speed. We found that cortex operates using a frequency-dependent transition between the latter two regimes **(Fig. 2g, Extended Data Fig. 3)**. At slow and infraslow frequencies (<1 Hz), coherence length spanned the entire cortex, matching the uniform medium model. At ∼1Hz, however, the coherence length dropped abruptly with a plateau at ∼2 mm, marking the emergence of propagation boundaries that compartmentalize the cortex. Above 12 Hz, a 1/f roll-off occurred, consistent with a scaling signature of the average local propagation speed (approximately 25 mm/s) notably much slower than the interhemispheric communication (2-5 mm/ms), suggesting distinct mechanisms for local versus interhemispheric communication.

To identify the underlying biology imposing these boundaries between 1–10 Hz, we applied non-negative matrix factorization (NMF)^32^ to the coherence matrix. This unbiased approach revealed five distinct dynamical structures remarkably consistent with the directional elements of Kochalka et al^33,34^: Anterolateral (AL), Active Sensory (AS), Medial Somatomotor (MS), Visual (V), and Medial Associative (MA); **Fig. 2h, Extended Data Fig 3d-e**]. Coherence length was significantly higher within these dynamical element networks than on average across the cortex (**Extended Data Fig 3d-e)**– with the exception of the Visual network, likely reflecting its known highly specialized and rigid retinotopic parcellation^35–37^ confirming that these borders place constraints on activity propagation. Furthermore, these boundaries partitioned the functional activity of the cortex significantly more accurately than did traditional cytoarchitectonic labels^38^ **(Fig. 2i)**, suggesting that the spatiotemporal logic of cortical dynamics is defined by this small number of discrete dynamical elements that govern the propagation of fast neural activity at the brain-spanning scale.

### Mesoscopic V-Ca linearity and state-dependent divergence

With these organizational principles of cortical electrical dynamics initially mapped, we next sought to determine how subthreshold fluctuations transition to suprathreshold spiking. By co-expressing the voltage indicator ASAP7y and the red-shifted Ca^2+^ indicator srGECO^39^ **(Extended Data Fig 4a-c)**, we achieved simultaneous dual-color imaging with CIM, capturing high-fidelity voltage signals alongside Ca^2+^ activity **(Fig. 3a-c, Extended Data Fig. 4d, Sup. Video 3)**. This approach allowed us to distinguish supra-threshold Ca^2+^-signals from sub– and supra-threshold voltage dynamics, with mesoscopic Ca^2+^-dynamics consistently lagging voltage dynamics by ∼60 ms **(Fig. 3c-d)**.

**Figure 3:**
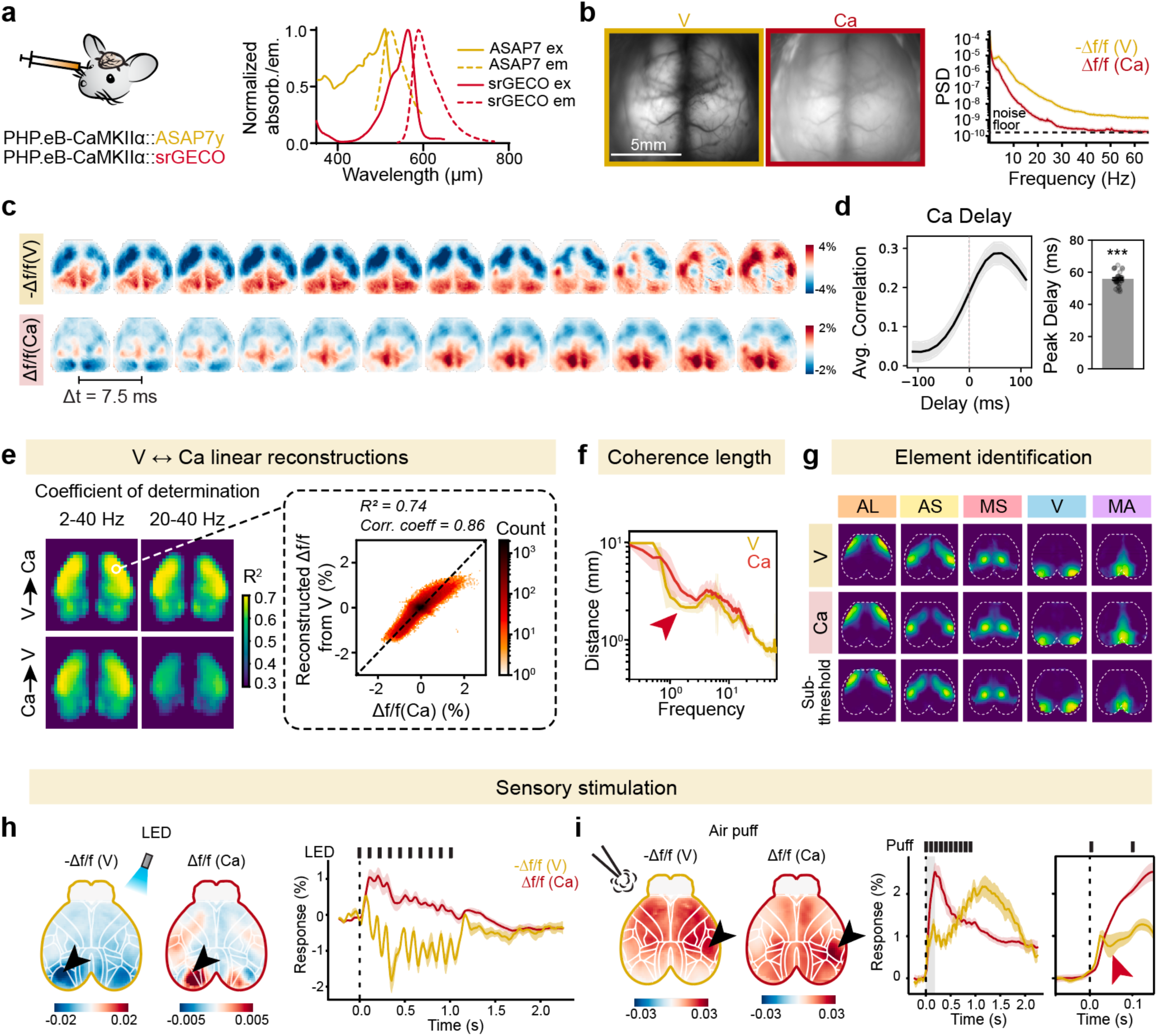
Simultaneous sub– and supra-threshold imaging reveals distinct dimensions of cortical dynamics. **a.** Dual-color imaging of voltage (V) and Ca^2+^ (Ca) dynamics (panels **a-d)**. Panel **a** shows co-expression strategy for dual-color imaging. *Left*: ASAP7y (V) and srGECO (Ca) are coexpressed by retroorbital injection of AAV-PHP.eB-CaMKIIα-ASAP7y and AAV-PHP.eB CaMKIIα::srGECO. *Right*: Corresponding optical excitation and emission spectra. **b.** *Left*: Representative wide-field images of simultaneously-captured V and Ca signals. *Right*: Power spectral density (PSD) of V and Ca signals. Noise floor is estimated by the frequency-independent floor of the PSD, reflecting shot-noise limit. **c.** Representative traces of simultaneously-acquired V and Ca activity (sampling rate, 133 Hz) show highly correlated activity, with a characteristic temporal delay in the Ca signal relative to V (7.5 ms between sequential imges; note similarity of 1^st^ V and 9^th^ Ca image (reflecting 60 ms delay). **d.** Correlation and temporal delay between V and Ca. *Left*: Average correlation across cortex against relative delay in Ca signal compared to V. *Right*: The maximum correlation is significantly delayed from 0 (p < 0.001, one-tailed t-test). Ca activity delayed by an average of 56ms from V; N = 4 mice, n = 32 sessions. **e.** Spatiotemporal transformations between V and Ca (panels **e-g**). Panel **e** shows predictive modeling of V-to-Ca and Ca-to-V transformations. *Left*: Mean coefficient of determination (R^2^) for linear (2-pole) bidirectional transforms across low (2–40 Hz) and high (20–40 Hz) frequency bands, averaged across N =4 mice, n= 16 sessions. *Right*: Measured Δf/f (Ca) signal versus the Ca trace predicted by a linear filter applied to negative Δf/f (V); representative session shown. The transform (for motor cortex ROI) exhibits an approximately linear relationship with a mild sigmoidal nonlinearity (see Extended Data Fig **3d**). **f.** Coherence length analysis of V and Ca signals. Both modalities indicate conserved functional barriers between networks; arrows denote a consistent coherence dip observed in both signals. Mean ± s.e.m.; N = 4 mice, n = 16 sessions. **g.** Elemental networks derived via coherence clustering. Parcellations are shown for V, Ca, and Subthreshold V (the residual voltage signal after regressing out the transformed Ca component), demonstrating conserved network boundaries across modalities. Representative data; similar results seen in N = 4 mice. AL: anterolateral; AS: active sensing. MS: medial somatomotor; V: visual; MA: medial associative. **h.** Sensory modulation of V and Ca activity (panels **h, i)**. Panel **h** shows visual stimulation evoked responses; arrows denote the primary visual cortex (V1) response. Trial-averaged activity (*right,* 10 Hz stimulus) shows divergence between V and Ca; note opposite sign and increased temporal richness of V measurement. Mean ± s.e.m., N = 3 mice. **i.** Air puff-evoked response. Trial-averaged activity (10 Hz whisker air puff) shows concurrent V and Ca increases in barrel cortex (arrows). *Right*: Fast-timescale initial suppression (arrow) observed only in V signal. Mean ± s.e.m., N = 3 mice.

We first quantified the shared information between these two signal streams. Coherence between voltage (V) and Ca^2+^ (Ca) genetically-encoded reporter signals was remarkably high from 2–20 Hz (**Extended Data Fig. 4e)**; using a second-order linear model, we predicted Ca from V with a coefficient of determination nearing 75% over this range **(Fig. 3e, Extended Data Fig. 4f)**. While higher frequency (20-40 Hz) V signals carried sufficient information to reconstruct Ca dynamics, the reverse was not possible at higher frequencies, where Ca signals lacked the necessary temporal information to recover fast voltage fluctuations. Analysis of reconstruction revealed that the mapping from V to Ca was nearly linear **(Fig 3e)**. Although this transformation will be inherently nonlinear at the single-cell level (governed by the threshold for activating voltage-gated Ca^2+^ channels), integrating thousands of neurons with varying activity levels within a mesoscopic pixel (∼130 x 130 µm) creates a broad linear operating regime **(Extended Data Fig. 4g)**. Consequently, Ca and V exhibit nearly identical coherence length scaling, and share the same elemental network parcellations **(Fig. 3f-g)**. Even the subthreshold residuals (isolated by subtracting the Ca^2+^-derived estimate from raw voltage) obeyed these elemental activity boundaries **(Fig. 3g)** which therefore appear to similarly constrain both subthreshold and suprathreshold dynamics.

This relationship between V and Ca shifted during spontaneous animal behavior. By tracking facial motion **(Extended Data Fig 5a-b),** we observed that while movement bouts triggered a global DC increase in both signals, fast fluctuating voltage power (>0.5 Hz) decreased across all frequency bands and networks, revealing a state transition from highly coherent, quiescent activity to a de-correlated state of arousal^40^ **(Extended Data Fig. 5c-e)**. In an intermediate frequency regime (0.5-9 Hz), V and Ca modulation diverged **(Extended Data Fig. 5d)**, and external stimuli further decoupled these signals. During 10 Hz visual flashes, V and Ca became anti-correlated, likely driven by a strong bilateral hyperpolarization in primary visual cortex that remained undetectable in the Ca channel **(Fig. 3h)**; this population-level separation may break the linear V to Ca transform by splitting the local voltage distribution into more-depolarized and more-hyperpolarized populations, a distinction invisible to more traditional Ca-based widefield imaging. Conversely, 10 Hz whisker puffs evoked concordant V and Ca responses in somatosensory cortex, though V measurement revealed far more temporally rich dynamics including a rapid, 10 ms suppression phase at the onset of the first puff that Ca indicators failed to capture **(Fig. 3i)**, as well as a late depolarization rising and continuing after termination of the stimulus train. Together, these results demonstrate that while the cortex operates linearly at the mesoscopic level between population subthreshold and suprathreshold activity during quiet wakefulness, behavior and sensory input can drive the system into regimes where these dynamics diverge, with far greater richness of information revealed by CIM voltage signal maps.

### Pharmacological perturbation of dynamical architecture

One of the major motivations of this approach is to gain insight into brainwide dynamics that may become maladaptive in mysterious brain states relevant to neuropsychiatry. We therefore next probed responsiveness of CIM-mapped dynamics to clinically-relevant perturbations, beginning with psychoactive compounds with distinct mechanisms of action, to test robustness of the patterns observed and to explore possible mechanisms of drug action. We began by administering powerful psychotropics representing distinct classes: ketamine, diazepam, or LSD in 15-minute sessions following saline control administration **(Fig. 4a-b).** Drug doses were selected for clinical relevance while maintaining consciousness in all cases (tracked with facial motion energy); the ketamine dose (administered at 90mg/kg) was chosen to be near-anaesthetic while maintaining consciousness befitting its approved clinical use in anesthesiology, while diazepam (anxiolytic dosing) was administered at 2mg/kg) and LSD (hallucinogenic dosing with no anesthetic properties) was administered at 0.3 mg/kg).

**Figure 4:**
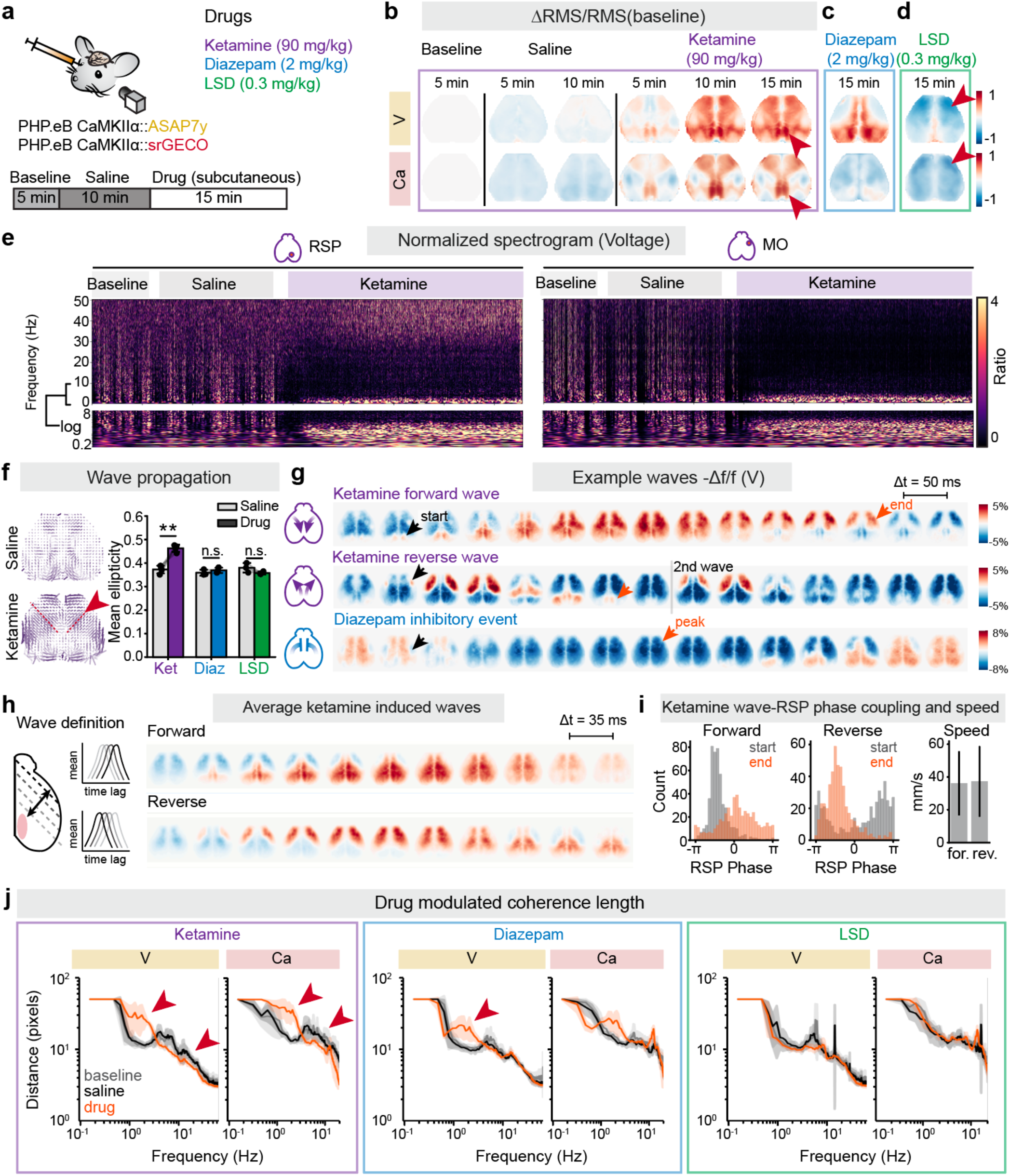
Pharmacological perturbation of activity propagation through elemental networks. **a.** Experimental strategy and pharmacological protocol. ASAP7y and srGECO are co-expressed for dual-modality imaging for voltage (V) and Ca^2+^ (Ca). Each session consists of a 5-min baseline, 10-min after saline administration control, and 15-min after drug administration. Ketamine (90 mg/kg), diazepam (2 mg/kg), and LSD (0.3 mg/kg) were administered subcutaneously. Only one drug was administered per 24-h period, with a minimum washout of 72 h between different pharmacological sessions. **b.** Psychotropic drug-induced shifts in cortical power (panels **b–d)**. Panel **b** shows power modulation over time following ketamine administration. Plots show mean changes in power (change in RMS power normalized to baseline RMS power) in 5 min increments. Ketamine at this dose induced widespread power in both V and Ca across the cortex, with the most pronounced shared effects in retrosplenial cortex (RSP; arrows); N =4 mice. **c.** Power modulation during the active phase of diazepam (10–15 min post-injection). In the V signal, power increased within medial associative and visual elements while decreasing elsewhere. Ca power largely failed to reflect these changes, although there was a detectable posteromedial increase along with a more general global decrease in other regions; mean changes shown across N = 3 mice. **d.** Power modulation during the active phase of LSD (10–15 min post-injection). Global power reduction was observed, predominantly within anterior cortical regions (arrows); mean changes shown across N = 3 mice. **e.** Spatial-temporal dynamics of drug induced activity (panels **e-g**). Panel **e** shows normalized spectrogram of RSP (*left*) and MO (*right*) cortex during ketamine administration. A sharp increase in δ power was observed corresponding to RSP oscillations in RSP and to a lesser extent in MO, along with a localized increase in γ power within RSP. **f.** Analysis of local wave propagation across pharmacological conditions. *Left*: Average map of local wave propagation during saline vs. ketamine (N = 4 mice). Color saturation represents ellipticity (indicating directional bias) and vector length inversely represents propagation speed (longer vectors indicating slower propagation). Arrow indicates a region with significant increase in wave propagation directionality and decrease in velocity under ketamine. *Right*: Mean ellipticity (directionality) changes under drug show significant increases in ketamine (p = 0.0086 paired t-test, N = 4 mice) but not in diazepam or LSD (N = 3 mice). **g.** Single-trial examples of cross-network phenomena. In ketamine, forward and reverse waves are shown with arrows denoting initiation (black) and termination (orange) points. In diazepam, arrows indicate the onset (black) and peak (orange) of a rapid, cortex-wide inhibitory event. **h.** Characteristics of ketamine-induced cortical waves (panels **h–j**). Panel **h** shows quantification of distributed cortical waves and relationship to RSP oscillations. *Left*: Wave identification based on threshold crossings across four parallel oblique axes (oriented 45° relative to midline, excluding RSP) in forward or reverse temporal order. *Right*: Trial-averaged profiles of identified forward and reverse waves. **i.** Ketamine wave-RSP coupling and propagation speed. *Left and middle*: Phase-locking of cortical waves to RSP oscillations. Distribution of initiation times for forward waves and termination times for reverse waves relative to the RSP oscillatory cycle (N = 4 mice). Both events are sharply phase-locked to the delta-band RSP oscillation (0.5–4 Hz); phase was estimated via Hilbert transform of the filtered RSP signal. *Right*: Propagation kinetics. Mean ± s.d. (N = 4 mice) of forward and reverse wave speeds. Velocities were calculated based on transit time between the first and last oblique axes defined in (**h**). Both wave types propagate at approximately 37 mm/s. **j.** Coherence length analysis across drug (orange) conditions. The coherence dip corresponding to elemental network boundaries is abolished by ketamine and reduced by diazepam, but remains intact under LSD. Average propagation speed is significantly reduced under ketamine. Mean ± s.e.m (N =4 for ketamine, N = 3 for diazepam and LSD).

These three interventions induced distinct shifts in CIM-measured neural activity, evidenced by changes in root-mean-square (RMS) fluctuations of V and Ca signals relative to baseline **(Fig. 4b-d, Extended Data Figs. 6a-c, 7a-c, 8a-c, Sup. Video 4)**. Consistently, V imaging with CIM revealed far greater complexity of brainwide dynamical maps (compared to Ca imaging with CIM) across all three classes of agent, including medial (LSD) and midline/posterior (diazepam) fluctuating dynamics not apparent with Ca imaging **(Fig. 4b,c)**. Consistent with previous reports at lower doses, ketamine elicited a band-limited power increase in the delta range (0.5–4 Hz) most prominent in retrosplenial cortex (RSP)^41–43^ **(Fig 4e, Extended Data Fig. 6a)**. In addition to this delta oscillation detected by CIM voltage imaging, we also observed with CIM voltage imaging, at this ketamine dose, a focal elevation of high-speed gamma power (30–60 Hz) within RSP that correlated with delta band activity **(Extended Data Fig. 6e)**, and which contrasted with a widespread gamma decrease in other cortical regions **(Fig. 4e, Extended Data Fig. 6a-b)**.

We next quantified how these agents might affect spatiotemporal propagation of electrical activity **(Fig. 4f, Extended Data 7e, 8d)**, leveraging the mapping capabilities of CIM. Among the compounds tested, ketamine induced the most fundamental reorganization, biasing propagation along a dominant axis spanning the posterior midline to anterolateral cortex with a significant increase in mean ellipticity, reflecting a transition toward highly axis-aligned wave propagation **(Fig. 4f)**. These changes were closely linked to the emergence of specific dynamic events: anterior-to-posterior (reverse) and posterior-to-anterior (forward) traveling waves **(Fig 4g-h, Extended Data Figs. 6d)**. The average wave speed was 37mm/s, remarkably consistent with the propagation velocities derived from our coherence length analysis in spontaneous activity without pharmacological intervention **(Fig 4i)**. Notably, these traveling waves (that were identified independent of RSP dynamics) were strongly correlated with RSP activity and phase-locked to 1–3 Hz oscillations previously linked to ketamine’s dissociative effects **(Fig 4h-i, Extended Data Fig 6e)**^41–43^; this rhythmic RSP phase relationship further correlated with mouse behavior **(Extended Data Fig 6f).** In contrast, diazepam and LSD did not induce such fundamental dynamical-map reorganization despite eliciting profound changes in neural activity. While diazepam triggered rapid, brain-wide hyperpolarization **(Fig 4g)** and LSD significantly reduced anterior cortical activity, neither agent significantly altered mean ellipticity (the established principal axes of propagation) (**Extended Data Figs 7, 8**).

Finally, we evaluated how these perturbations might alter the frequency-dependent scaling of coherence length **(Fig. 4j)**. At these doses of ketamine or diazepam but not LSD, the characteristic dip in coherence length (see **Figure 2**) marking the emergence of network boundaries at ∼1Hz, was selectively blocked; we also were able to measure a shift, that was selective to ketamine, in the 1/f roll-off associated with local propagation shifting to lower frequencies, indicating a measurable reduction in local conduction velocity. Interestingly, though the average conduction velocity dropped detectably in ketamine, the conduction velocity of the induced large-scale waves (37mm/s) remained slightly above spontaneous baseline. Under LSD however, despite the extraordinary perturbation to local power statistics of neural activity **(Fig 4d)**, cortex-wide coherence structure remained largely intact **(Fig. 4j)**. Together, these results demonstrate that different classes of psychotropic agent interventions distinctly and precisely reconfigure the spatiotemporal rules of cortex-spanning communication, with individual signatures measurable with CIM-based imaging.

### Casually testing dynamic architecture with optogenetic and human genetics-guided interventions

While pharmacological interventions provide substantial basic and clinical relevance for CIM-based dynamical mapping, optogenetic interventions provide far more spatiotemporally precise perturbation capability and thus may offer greater insight into the precise causal mechanisms of these dynamical patterns. Therefore, we co-expressed the green light-activated GEVI ASAP7y alongside rsChRmine (a designed red-actuated high-sensitivity trimeric pump-like channelrhodopsin^44^), with both probes under CaMKIIα promoter control **(Fig. 5a, Extended Data Fig 9a-c)**. To achieve precise, high-intensity optogenetic stimulation at arbitrary locations, we integrated a two-axis microelectromechanical (MEMS) mirror to steer a focused laser beam (200 μm spot size) **(Fig. 5b-c)**. Although the designed rsChRmine spectrum still partially overlaps with the excitation wavelength of ASAP7y, we found that the high collection efficiency of CIM platform permitted imaging at power densities orders of magnitude below the threshold for incidental opsin activation **(Fig. 5c, Extended Data Fig. 8d)**.

**Figure 5:**
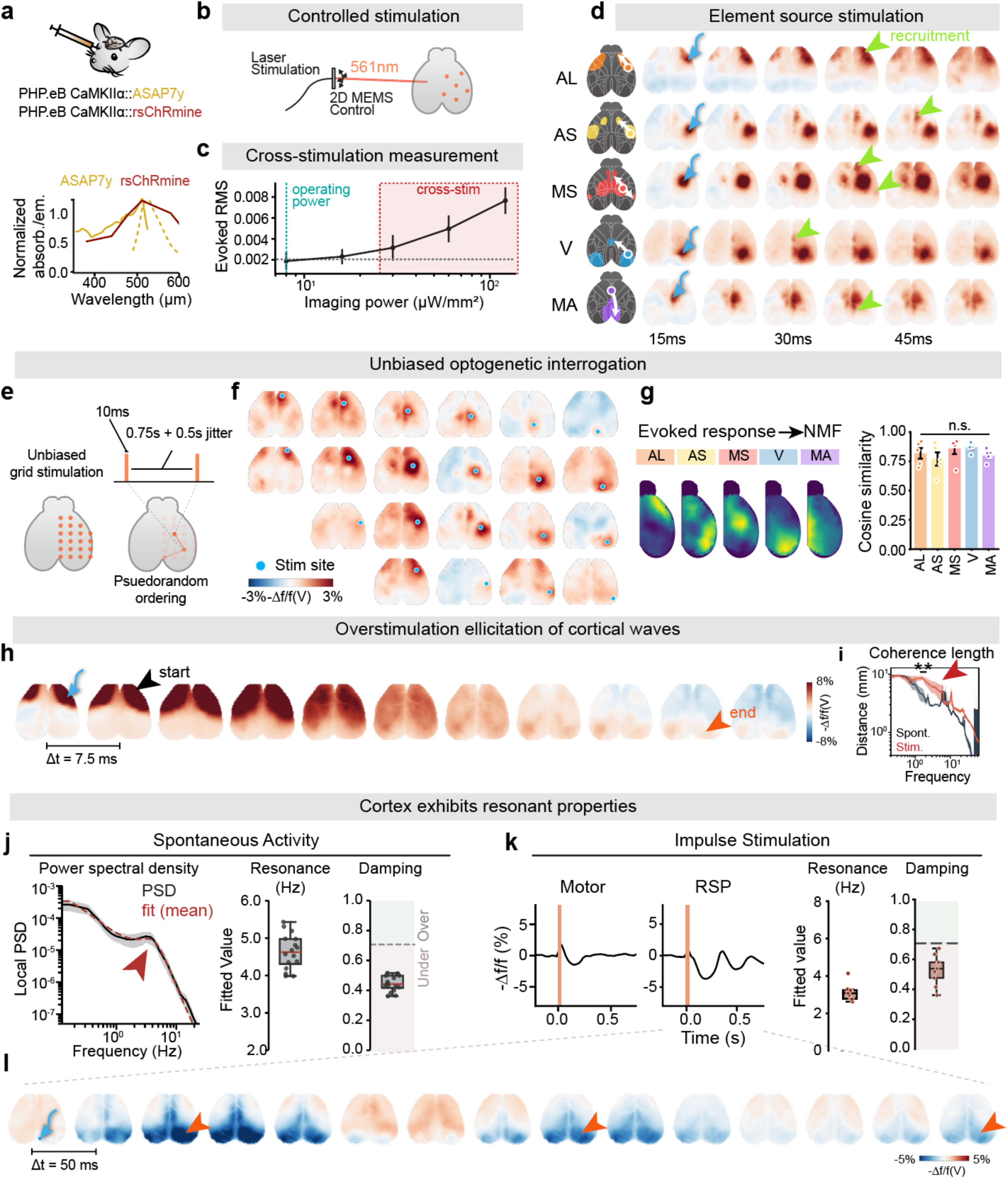
Precise causal tests of cortical organization principles. **a.** Integration of voltage imaging and optogenetic control (panels **a-c**). Panel **a** shows expression strategy and spectral characterization of the relevant tools. *Top:* ASAP7y (voltage sensing) and rsChRmine (optogenetic control) were co-expressed via retro-orbital injection of AAV-PHP.eB CaMKIIα::ASAP7y and AAV-PHP.eB CaMKIIα::rsChRmine. *Bottom:* Excitation (solid line) and emission (dotted line) spectra for ASAP7y superimposed on action spectrum of rsChRmine (red line). **b.** Precision optogenetic targeting. Stimulation was controlled via a micro-electromechanical system (MEMS) for high-resolution 2D targeting across the entire cortical field. **c.** Cross-stimulation measurement showing steady and then increasing RMS activity (measured in the stimulated area) during a 0.5s long stimulation. Stimulus-induced artifacts were negligible at powers <15 μW/mm^2^; subsequent experiments (below) therefore utilized 8 μW/mm^2^ operating power. **d.** Optogenetic stimulation at source locations recruited full specific network elements (panels **d-g**). Panel **d** shows targeted stimulation of specific element sources. Representative trial-averaged trace (similar results in N = 6 mice) shows that activation of the source within a network element recruited entire functional element; blue arrows indicate specific stimulation sites, and green arrowheads denote recruitment of non-local sites across the dynamic elements**. e.** Unbiased grid stimulation paradigm. Random-walk stimulation across a cortical grid (10-ms pulses, 0.75-s interval ± 0.5-s jitter) was used to map evoked responses. **f.** Representative evoked response. Trial-averaged cortical activity (all 45 ms post-pulse from a single animal; similar results across N = 6 mice) following typical moderate optogenetic stimulation (10ms, 4 mW/mm^2^ calculated cortical irradiance; Methods). Blue dots indicate specific stimulation sites. **g**. Functional parcellation of evoked activity. *Left*: Non-negative matrix factorization (NMF) of responses decomposed cortical activity into the same elemental networks identified during spontaneous activity. *Right*: Cosine similarity between evoked recruitment patterns and established functional parcellations; N = 5 mice. No significant differences were noted in recruitment among elements; statistics determined by one-way ANOVA (p = 0.1097). **h.** Overstimulation (5x, to 20 mW/mm^2^ calculated cortical irradiance; Methods) overcame network barriers to induce global waves (panels **h, i)**. Panel **h** shows induction of global cortical wave activity recruiting large, cross-element domains; blue arrow denotes stimulation site and arrowheads mark start and end of the cortical wave. Trial average of a representative mouse from N=4 mice. See extended figure 9 and supplementary video 5. **i.** Overstimulation abolishes coherence-length dip (arrow), indicating a blurring of element boundaries; coherence length differs significantly between groups in this 1.4-2Hz window (p = 0.008, two-tailed Welch’s t-test; mean ± s.e.m., N = 4 mice for spontaneous activity, N = 5 mice for overstimulation). **j.** Cortex as an under-damped resonant medium (panels **j–l**). Panel **j** shows spectral analysis of spontaneous activity. *Left*: Pixel-wise power spectral density (PSD) reveals a characteristic resonance peak (arrow) (N = 4 mice, n = 25 sessions); dashed lines indicate fits using median resonance frequency and damping coefficients. *Middle, right*: Distribution of fitted resonance frequencies and damping coefficients; damping values fall within the underdamped regime (< 1/√2). **k.** Kinetics of overstimulation-induced local response. *Left*: Trial-averaged evoked responses (in motor cortex and RSP) display damped oscillations, RSP shows a more prolonged oscillatory decay. *Middle, right*: Distribution of fitted resonance and damping coefficients averaged across 21 stimulation sites (N = 5 mice, n = 9 sessions). **l.** Underdamped RSP oscillation. Trial averaged whole-cortex dynamics of a representative RSP stimulation show oscillatory network recruitment at ∼3Hz. Orange arrow indicates stimulation site; orange arrowheads highlight successive oscillatory hyperpolarizations.

We optogenetically stimulated the “source” nodes of each of the five directional elements of cortical dynamics that we previously identified^33,34^ (**Fig. 2h, 3g**). This spatiotemporally-specific stimulation in each case was found to faithfully recruit the corresponding full directional element from source to sink (largely ipsilaterally, with a reduced contralateral contribution; **Fig. 5d)**, revealing that the functional boundaries identified from the spontaneous activity of quiet wakefulness **(Fig. 2)** in fact causally govern the spread of evoked cortical neural activity. To further probe the causality of spatial dynamics, we performed an unbiased optical/computational screen with interrogation at 21 hemispheric grid locations tiling dorsal cortex **(Fig. 5e-g)**. Delivering moderate (10ms/4mW at source; mean ∼4 mW/mm^2^ calculated across cortical thickness^45^) optogenetic stimuli was found to recruit well-structured local and non-local activity **(Fig. 5f, Sup. Video 5)** that, when decomposed via NMF, recovered the five naturally-occurring directional elements observed in spontaneous waking activity **(Fig 5g)**.

The optogenetic stimulus intensities used here were by design comparable to those typically used and recommended for optogenetics^46–51^, and the resulting observation of naturalistic patterns is consistent with prior work demonstrating that targeted optogenetic stimulation gives rise to naturalistic dynamics at the level of local ensembles, regional circuits, and brainwide activity^52,53^. However, in some cases it is experimentally of interest to elicit stronger or longer activity patterns than would naturally occur, to probe the boundary conditions of natural activity and to generate intentionally perturbative or even lesion-like effects^54,55^. We therefore sought to explore this dimension afforded by the flexibility of CIM integrated with precision optogenetics. We found that overstimulation (mean ∼20 mW/mm^2^) in this way could break the naturalistic boundary conditions, evoking more broadly-distributed activity patterns **(Extended Data Fig. 9e; Sup. Video 5)** that crossed natural functional borders and launched reverse or forward traveling waves; as a result, the characteristic dip in coherence length was abolished **(Fig. 5h-i, Extended Data Fig 9g-h)** as also seen in a subset of specific pharmacological interventions **(Fig. 4)**. Despite the elicitation of waves and loss of spatial compartmentalization, NMF decomposition of evoked signals could still largely recover the elemental network motifs **(Extended Data Fig. 9f)**, suggesting that overstimulation induces new wave dynamics without breaking the fundamental structure governing cortical activity.

Surprisingly, we observed that high-intensity cross-element cortical waves were sometimes followed by a secondary “rebound” wave appearing hundreds of milliseconds after the initial stimulus (**Extended Data Fig. 9g**). The phenomenon suggested that the cortex might have the capability to function as a resonant medium. To investigate this possibility, we first examined the local power PSD in spontaneous activity recordings. While the PSD followed the expected scale-invariant 1/f^-α^ distribution at high frequencies, we observed a distinct spectral peak between 3–5 Hz that was consistent across mice and cortical areas, present in both Ca and V recordings **(Fig 5j, Extended Data Fig. 9j-k)**. While it is not uncommon to find such peaks in electrical recordings^56,57^, the full spectrum—specifically the peak location and 1/f^-2^ roll-off—together strikingly resembled the response of an underdamped oscillator (**Extended Data Fig. 9i**). In the time domain, as the damping of an oscillator drops below the critical threshold (1/√2), the dynamics from an impulse transition from direct exponential decay to oscillatory ringing where the system repeatedly overshoots its equilibrium. To test if this spectrum did indeed reveal this resonant property, we examined temporal responses to brief optogenetic impulses. High-intensity stimulation (15ms at mean ∼20 mW/mm^2^) elicited prolonged 2–3 Hz ringing across cortex, particularly in posterior cortical regions (RSP and visual) **(Fig. 5k-l, Extended Data Fig. 9l-m)**, precisely as predicted for an underdamped resonator.

This low-frequency oscillatory signature resembled ketamine-induced dynamics **(Fig. 4),** suggesting that cortical resonance is an intrinsic tissue property that can be unmasked by specific pharmacology (ketamine) or intense stimulation. Such resonance may reflect the aggregated behavior of multiple mechanisms; for example at the cellular level, HCN channel-mediated I_h_ currents confer intrinsic resonance properties on individual neurons in the 4–10 Hz range^58–61^, while at the circuit level excitatory-inhibitory recurrence can sustain oscillatory activity through feedback inhibition^62–66^, and at the brainwide level thalamocortical loops– intrinsically capable of generating delta and spindle rhythm– can provide long range pathways that both generate and synchronize low-frequency rhythms across cortex^67–70^. How these disparate mechanisms (each operating at distinct spatiotemporal scales) may converge to produce coherent band-limited brain-wide resonance remains to be determined, in exploration which may be enabled by the CIM framework for capturing emergence and propagation of resonant dynamics with suitable scope and spatiotemporal resolution. Together, these results provide causal evidence that cortex acts as a resonant medium with modular boundaries, and that these boundaries (while governing orderly and perturbation-resistant performance under typical conditions) can be overcome in specific conditions, thereby reshaping the spatiotemporal rules of high-speed electrical activity propagation.

Finally, we asked if CIM-based voltage-mapping of cortical spatiotemporal dynamics would reveal measurable changes caused by human disease-relevant genetic perturbation. We began by exploring cortical dynamics in the widely-studied *Cntnap2* mouse model derived from autism spectrum disorder (ASD) genertics^71,72^, by expressing ASAP7y for CIM imaging in littermate-matched *Cntnap2* knockout (KO) and wild-type (WT) mice. Although *Cntnap2*^KO^ mice exhibit abnormalities in social and stereotyped behaviors, and have been reported to show reduced density of inhibitory interneurons in cortex along with elevated excitation-inhibition balance^72–74^, when mapping spontaneous activity we observed no significant differences in properties of the cortex-spanning directional elements **(Extended Data Fig. 10a-b)**, local wave structure **(Extended Data Fig. 10c)**, or power distribution in either voltage or Ca^2+^ (**Extended Data Fig. 10d**). While these results suggested that the modular dynamical organization of the dorsal cortex had remained largely intact in knockout animals, clear element-specific changes emerged in the context of external sensory stimuli. Airpuff-evoked voltage dynamics unveiled a significant change in adaptive post-depolarization inhibition in *Cntnap2*^KO^ mice, specifically localized to the active sensory (AS) dynamical element (75% cosine similarity; **Extended Data Fig. 10e-g)**. This specific phenotype, likely related to dysregulated excitatory-inhibitory balance characteristic of *Cntnap2*^KO^ animals, may bear relevance to human ASD wherein impaired adaptation to ongoing sensory stimuli represents a core contributor to clinically-significant distress and dysfunction. Moreover, this dysfunction, masked during spontaneous activity but emergent during evoked fast electrical signaling, could not have been readily resolved without the CIM voltage sensing framework, suggesting potential utility of this approach for identifying circuit-level signatures of adaptive dynamics as well as maladaptive states related to neuropsychiatric disorders.

## Conclusions

Neocortex transmits and integrates information across distributed neuronal networks spanning the mammalian brain. While local features of neocortical architecture are conserved across regions and species, overarching principles underlying the high-speed and long-range coordination of neural activity have remained largely mysterious. By developing conformal immersion microscopy (CIM) and integrating CIM with high-performance genetically-encoded voltage indicators, we were able to reveal spatial and temporal rules that govern activity propagation across the mouse dorsal cortex.

CIM solves two optical problems that have constrained mesoscopic activity imaging: insufficient photon collection and inability to maintain focus across the curved cortical surface. Speed-boosting focal-length reducers and a custom field-flattening lens address these jointly, yielding the signal-to-noise necessary to resolve single-trial dynamics at millisecond timescales without photobleaching — an enabling combination for the broader class of next-generation genetically encoded fluorescent indicators. Critically, both advances are implemented with accessible components, making CIM readily adoptable into existing mesoscopic systems. As implemented here, CIM operates at a spatial scale optimized for network-level imaging and cannot resolve single-cell dynamics, constraining mechanistic interpretation to the mesoscale. Extending CIM toward diffraction-limited resolution would bridge network-level phenomena to their cellular substrates, providing an “atomic” or component-level description of the emergent properties elucidated herein.

The advances provided by CIM in whole-cortex microscopy in sensitivity and coverage combine well with recent improvements in voltage imaging exemplified by ASAP7y. We demonstrate that this combination enables direct measurements of electrical dynamics spanning both subthreshold integration and suprathreshold output across infra-slow to high-gamma frequencies, in a way that calcium imaging, constrained by the ∼100-ms extrusion rates for calcium^75^, fundamentally cannot. In combination with physics-inspired analysis, CIM revealed the timescales and spatial organization of hemispheric and local communication, and parcellated the cortex into discrete functional elements — with voltage imaging further revealing that even the subthreshold component of voltage, isolated by subtracting the calcium-predicted signal, follows the same functional boundaries, suggesting that these elements constrain both integration and output. Beyond network organization, voltage imaging exposed fast subthreshold events invisible to calcium: rapid hyperpolarization evoked by visual stimuli, and a 10ms suppression at whisker puff onset. The ability to differentiate subthreshold integration from suprathreshold output at scale opens new avenues for studying how top-down neuromodulation and bottom-up sensory inputs are integrated within specific functional territories. The resolution of high-gamma dynamics cortex-wide further opens the largely unexplored question of how fast oscillations — including sharp-wave ripples — propagate and coordinate across the neocortical surface.

Perturbing cortical dynamics with psychotropic drugs (here ketamine, diazepam, and LSD) revealed that the spatiotemporal architecture identified during quiet wakefulness is largely robust to even high-dose pharmacological challenge, yet voltage imaging revealed substantially richer dynamics than calcium alone could provide, including elevated focal RSP gamma activity under ketamine and midline fluctuation under LSD and diazepam. The ketamine intervention in particular disrupted the coherent-length signature of functional element boundaries, giving rise to cortex-spanning travelling waves; the LSD intervention, despite dramatically altering neural activity, left coherence length intact. These signatures suggest CIM is well positioned to uncover how psychotropic drugs reconfigure the spatiotemporal rules of cortical dynamics. A systematic screen across compounds and doses may reveal conserved classes of neural dynamics underlying altered states of consciousness and cognitive function.

Optogenetic and genetic interventions further demonstrated that this approach to voltage imaging provides causal and disease-relevant insight unavailable from calcium readout alone. Targeted stimulation confirmed that evoked propagation followed the same element parcellation identified in spontaneous activity; overstimulation launched cortex-wide traveling waves that, rather than abolishing elemental structure, appeared to sequentially recruit the underlying directional elements. The blurring of functional boundaries under pharmacological perturbation and the causal unmasking of resonant properties via optogenetics further characterized the cortex as a tunable, resonant medium. This physical framework, now established with high spatiotemporal precision and coverage in mice, may be extended to human participants using the more limited but well-established clinical tools for recording and stimulation, with diverse potential implications for bulk clinical manipulation of neural networks.

By identifying element-specific deficits in a model of autism, we demonstrate that this framework may be powerful enough to reveal mechanistic insights into mesoscopic disruptions of neural circuits. In the Cntnap2-KO autism model, spontaneous activity appeared intact in both voltage and calcium recording, yet evoked voltage dynamics unveiled an element-specific deficit in adaptive post-depolarization inhibition following sensory stimulation — a fast subthreshold phenotype invisible to calcium and absent from spontaneous recordings. Though limited here to a single genetic model under one sensory modality, these findings suggest that systematic application across disease models and behavioral contexts might uncover voltage-specific signatures of circuit dysfunction that remain invisible to conventional imaging approaches.

Going forward, this experimental and conceptual platform offers a scalable template for mapping dynamics across cells, circuits, and systems spanning diverse behaviors, genetic backgrounds, and disordered states. As genetically encoded indicators continue to evolve in speed and sensitivity, combining this framework with cell-type-specific targeting may allow for a granular deconstruction of the rules governing cortical communication, ultimately providing a foundation for identifying the precise electrical fingerprints of cognitive states and their dysregulation in neuropsychiatric disease.

## Acknowledgements

We thank S. Vesuna, S. Quirin, T.X. Liu, and all members of the Deisseroth lab, as well as I. Landau, M.Bilokur, H.S. Hunt, and all members of the Ganguli lab for input on the project and feedback on the manuscript. This work was supported by the National Institutes of Health grant U19NS118284 (K.D.), National Institutes of Health grant P50DA042012 (to K.D.), National Institutes of Health grant R01 MH086373 (K.D.), the Gatsby Foundation (K.D.), a Knight Initiative for Brain Resilience Catalyst Award (K.D.), the Keck Foundation (K.D.), R01 EY022638 (to T.R.C.), P30EY026877 (to T.R.C.), The Chan Zuckerburg Biohub, San Francisco (to T.R.C.), a Stanford Bio-X Seed Grant (to T.R.C.). UM1MH136462 (to M.Z.L.) and 1RM1NS132981 (to M.Z.L.). Y.W. is a Howard Hughes Medical Institute Fellow of The Jane Coffin Childs Memorial Fund. Y.A.H. was supported by the Stanford BioX Bowes Fellowship.

## Author Contributions

A.D.W., Y.W., J.K., and K.D. designed the study. A.D.W. developed CIM. Y.W., A.D.W., C.L., M.H., J.K., W.W., A.M., B.E.G., and C.K. performed animal experiments. Y.A.H., T.R.C., and M.Z.L. developed ASAP7y. N.A. and K.E.P. produced viral vectors. A.D.W., Y.W., J.K., and K.D. analyzed the experimental data, with assistance from S.G. and T.R.C. in analysis conceptualization. A.D.W., Y.W., J.K., and K.D. wrote the manuscript with input from all authors. K.D. supervised all aspects of the project.

## Data Availability

Processed data files, including high speed neural recordings and corresponding motion energy for all analysis, will be available as a repository on Zenodo.

## Code and Data Availability

All code used for data analysis and figure generation are available by repository. The widefield imaging datasets and processed connectivity matrices generated during this study will be made available along with deposition of custom Python scripts for preprocessing and computation in a public repository upon publication.

**Extended Data Figure 1:**
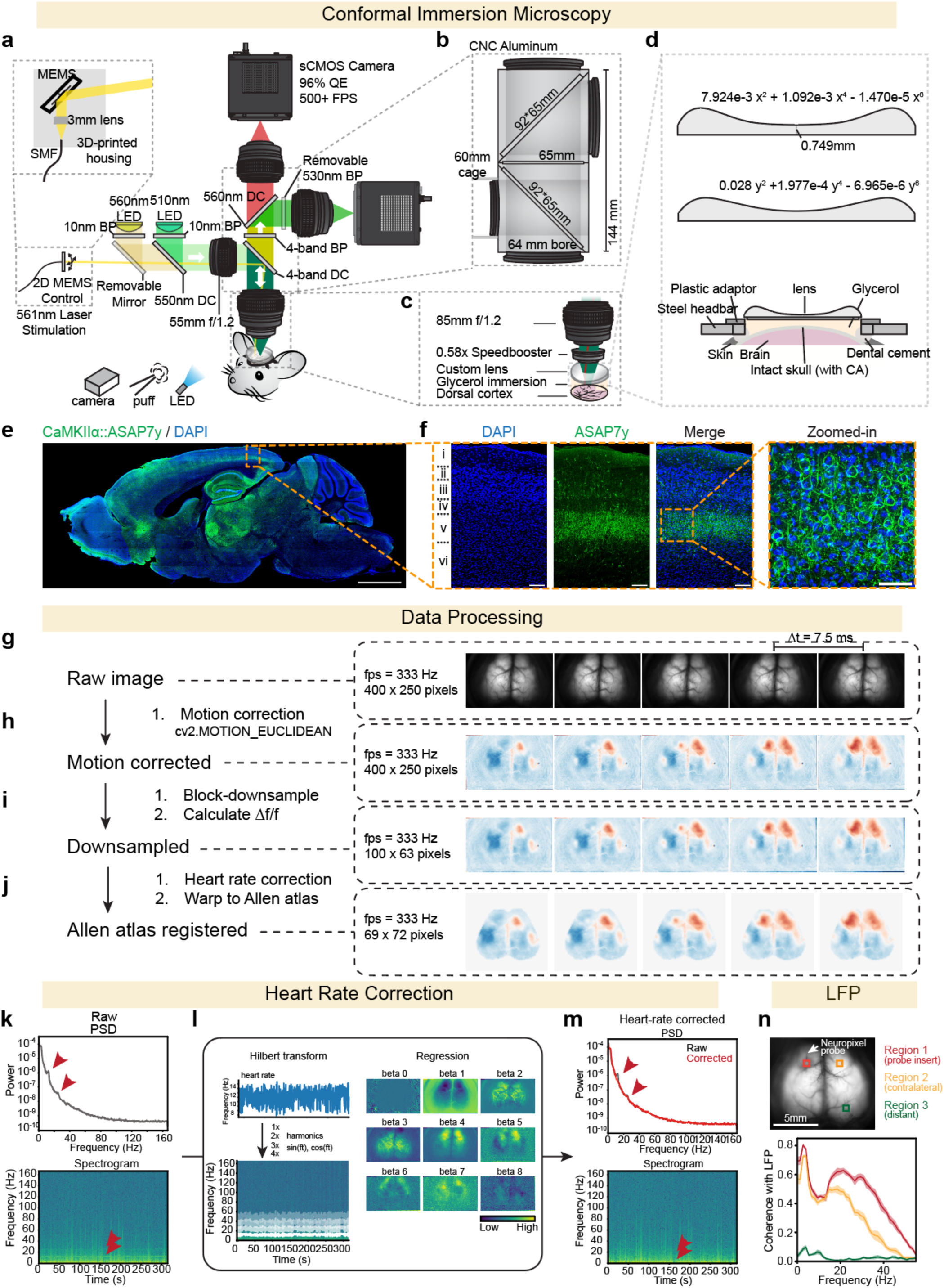
Conformal immersion microscopy and processing. **a.** Panels **a-d** show CIM design and construction; panel **a** shows full CIM microscope schematic. **b.** Core microscope housing is CNC machined from 7075 aluminum. Custom housing allows for larger filters to accommodate the large aperture with minimum (144 mm) throw. As the illumination path has less critical throw, it was constructed using off-the-shelf 60mm cage system parts. **c.** Objective lens is made up of f/1.2 85mm lens, custom printed lens (d), and is glycerol-immersion. **d.** Schematic of lens dimensions and mouse preparation. Lens is constructed with an x and y polynomial described by the equations here. Lens is printed with WaterShed XC 11122 resin and polished to a transparent finish. **e.** Representative sagittal section of CaMKIIαASAP7y; scale bar: 2mm**. f.** Representative images of ASAP7y expression in cortex (scale bar: 100 um) and zoomed-in view of layer V (scale bar: 50 um)**. g-j, g.** Data acquisition and pre-processing: raw 3200*2000 pixel images are captured from the Kinetix cameras and binned 8×8 with averaging to reduce to a size of 400*250 pixels before storage to save storage space. **h.** Images are motion corrected and the Δf/f is calculated. **i.** To further increase SNR, Δf/f is 4×4 downsampled to 100*63 pixels. **j.** Heart-rate correction and registration (warping) to the Allen Brain Atlas (69 × 72 pixels). **k.** Panels **k-m** show heart rate correction; panel **k** shows hemoglobin absorption at yellow wavelengths introducing noise at the heartbeat frequency and its harmonics (red arrows). **l.** Regression of heart-rate artifacts. The brain-wide average signal is bandpass filtered (9-13 Hz) to isolate the heartrate. A Hilbert transform extracts the time-dependent phase, and the first four harmonics (including sine and cosine components for phase-shift compensation) are regressed pixel-wise via linear regression. Maps show regression coefficients for a representative trace. **m.** The regressed signal exhibits significantly reduced noise at heartbeat-associated frequencies (red arrows). By maintaining constant regression amplitudes over time, local time-varying neuronal signals are preserved. **n.** Coherence between LFP and Δf/f fluorescence of ASAP7y across three ROIs. Plots show mean coherence across 10 1-minute periods, and shading shows s.e.m.

**Extended Data Figure 2:**
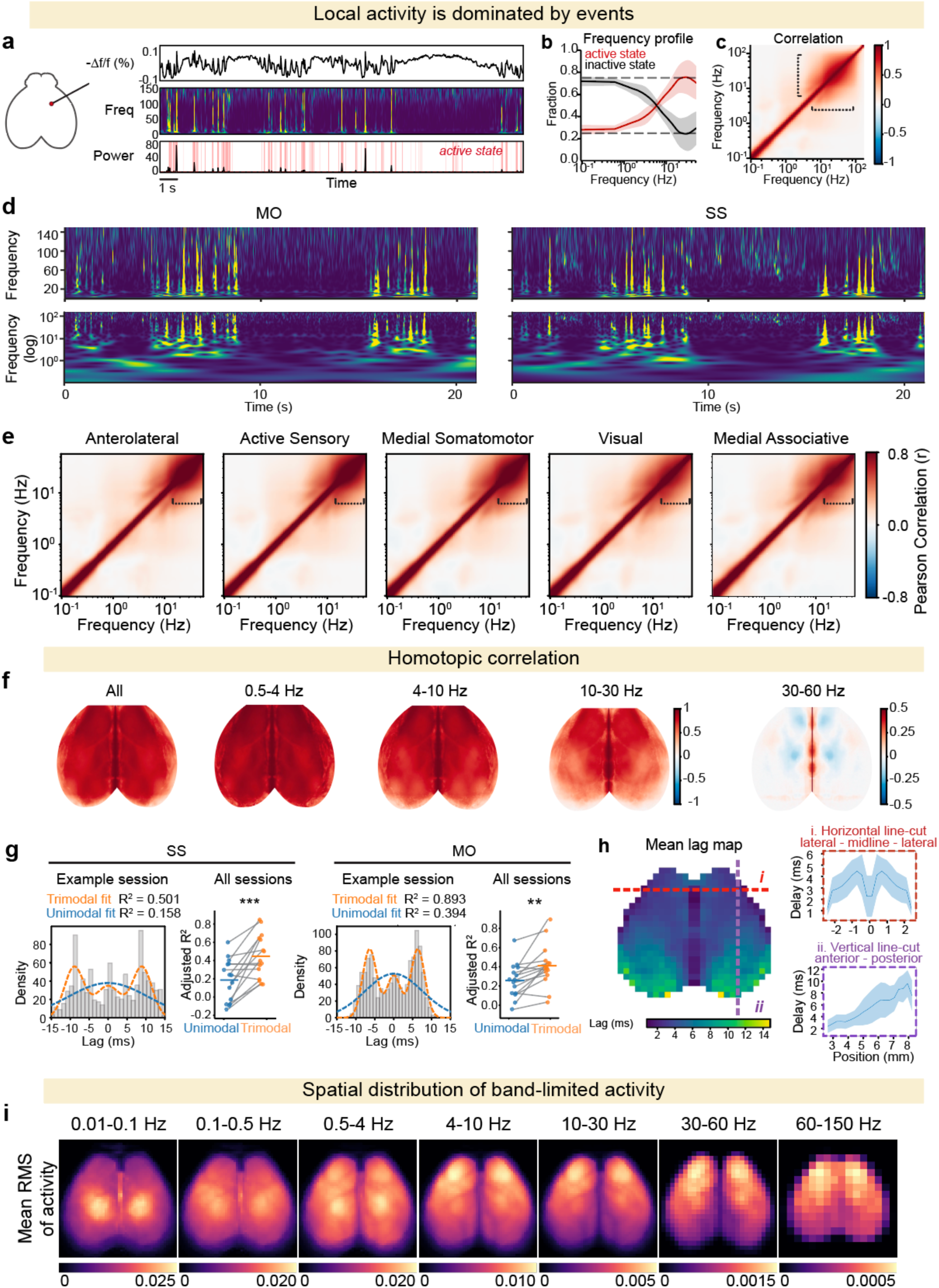
Spanning fast and slow timescales: emergent spatiotemporal structure of local cortical dynamics. **a.** Panels **a-e** show spontaneous local activity over long timescales is dominated by phasic events; panel **a** shows representative trace and spectrogram from an ROI in motor cortex, revealing banding in time. The bottom graph shows temporal evolution of active states of the ROI shown at left, defined as time bins wherein the average whitened power is in the top 25%. **b.** Fraction of power per frequency in the active (top 25%) and inactive (bottom 75%) states, averaged over all pixels. While low frequencies are more uniformly distributed, high frequencies are skewed towards the active state. Mean ± s.e.m.; N=4 mice, n=25 sessions. **c.** Summary graph of correlation between frequencies. To assess the intrinsic spectral organization of spontaneous cortical activity, time-resolved power spectra were estimated using a Morlet wavelet transform across a logarithmic frequency range (0.1 Hz to Nyquist). Pearson correlation coefficients were computed between all frequency pairs across valid pixels within the cortical mask and averaged across pixels, then across animals. Off-diagonal structure (dashed brackets) indicates coordinated power fluctuations across distinct frequency bands, suggesting multi-scale neural dynamics. The average frequency-frequency correlations showing that these higher frequencies that are skewed towards events are also highly correlated with each-other. Average data from N=4 mice, n=14 sessions. **d.** Exemplar whitened spectrograms of activity in motor and somatosensory areas showing bursts of activity in high frequencies and slower variation in low frequencies. **e.** Frequency-frequency correlations within network elements. For each pixel belonging to a given functional motif, time-resolved power spectra were estimated via Morlet wavelet transform across a logarithmic frequency range, and Pearson correlation coefficients were computed between all frequency pairs. Matrices were averaged across pixels within each motif, then across animals. Correlation structure is largely preserved across motifs, but high-frequency coupling (brackets) differs between motif types. Average data from N=4 mice, n=25 sessions **f.** Temporal complexity of homotopic correlations. Band-limited maps of homotopic correlations revealing reduced correlation with increasing frequency (example session shown; consistent across N=4 mice, n = 25 sessions) **g.** Cross-correlation peak time histograms reveal multi-modal delay distributions in spontaneous bilateral activity for somatosensory (SS) and motor (MO) cortices. Orange and blue dashed curves show trimodal and unimodal Gaussian fits respectively. Right panels compare adjusted R² across N=4 mice and n=25 sessions, demonstrating that the trimodal model significantly outperforms the unimodal model in both regions (*** p<0.001, ** p<0.01, paired two tailed t-test). **h.** The frequency-dependent reduction in correlation as shown in **f-g** indicated the possibility of a homotopic correlation delay. A measurement of this delay across the cortical surface is shown here in **h**. Line-cuts (red: horizontal, lateral-midline-lateral; purple: vertical, anterior to posterior) across this map show that delay does not scale linearly with distance and remains consistent across mice, with shading indicating standard deviation (N=4 mice, n=25 sessions). **i.** Spatial distribution of band-limited cortical activity. Mean RMS amplitude maps across seven frequency bands (0.01–150 Hz) derived from widefield voltage imaging. Each panel shows the spatial distribution of band-limited power averaged across pixels, sessions, and animals (N=4 mice, n=25 sessions). Note the progressive shift in spatial topography across bands: Low frequency (0.01-4Hz) shows roadly distributed bilateral activity, while high frequencies (4-150 Hz) highlight more anterior activities.

**Extended Data Figure 3:**
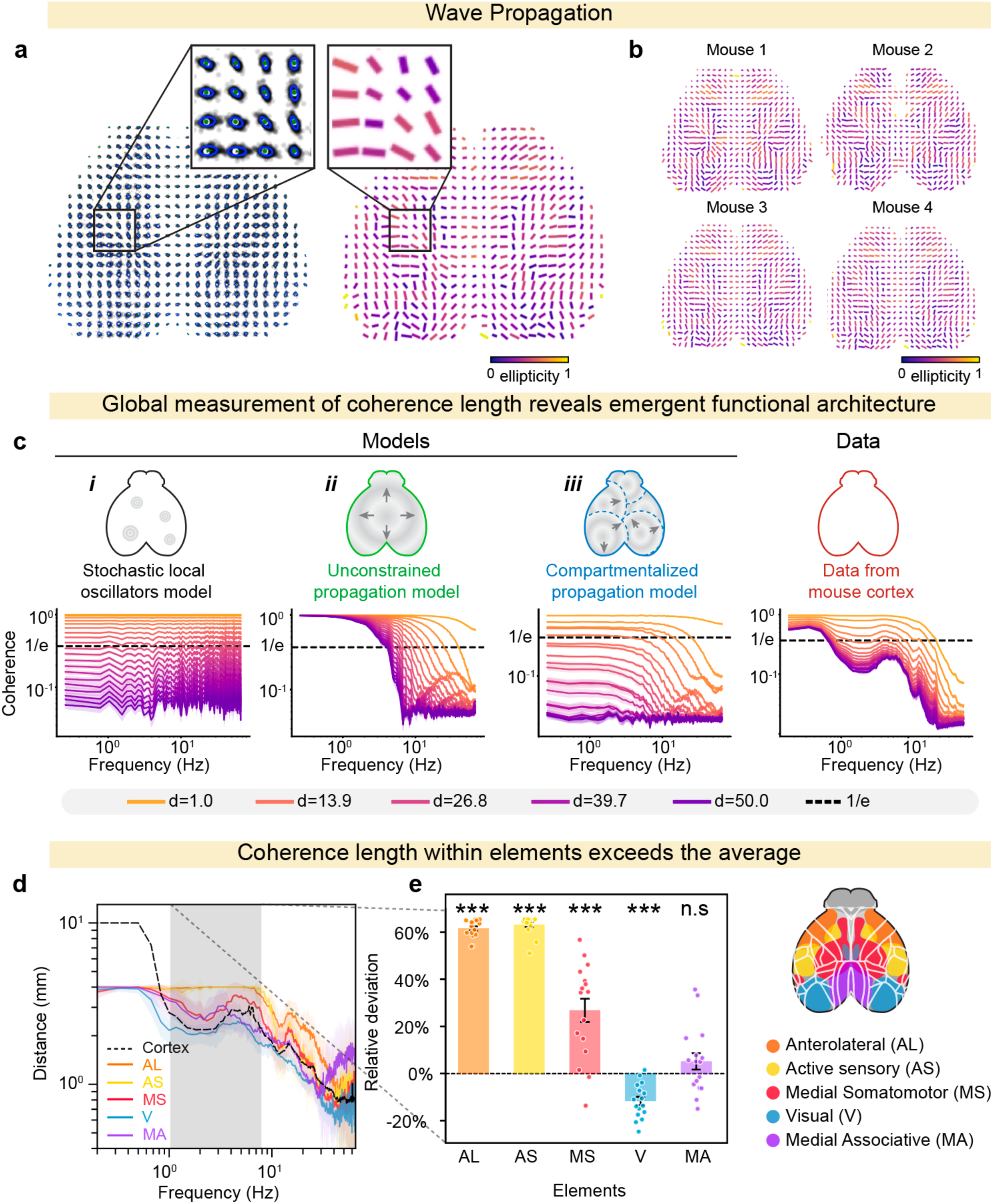
Emergent spatiotemporal structure of global cortical dynamics. **a.** Wave propagation map: delay vector map and corresponding propagation map from Fig 2e shown in more detail with zoomed in view. Higher ellipticity means stronger directional dominance. **b.** Propagation map of N=4 mice. **c.** Coherence as a function of frequency across pairwise distances comparing simple models versus real CIM recorded cortical data. Each line represents the coherence spectrum at a given pixel separation distance (d = 1.0–50.0 mm, orange to purple). Coherence length at each frequency is defined as the furthest distance whose coherence remains above the 1/e threshold (dashed line), with linear interpolation between discrete distance steps. The resulting coherence length as a function of frequency is summarized in Figure 2g. Three toy models are shown on the left: (i) stochastic local oscillators show distance dependent coherence decay that is uniform across frequencies, (ii) unconstrained propagation show high coherence at low frequencies and drop with distance at high frequencies, and (iii) compartmentalized propagation show frequency dependent coherence decay with increasing distance. Rightmost panel shows widefield voltage imaging data from mouse cortex, which have high coherence at low frequencies and frequency-dependent coherence decay at high frequencies. **d-e.** Coherence length vs. frequency within the dynamical elements. **d.** Characterization of coherence length vs. frequency inside element parcels, compared to cortical average. Mean ± s.e.m. from N = 4 mice, n = 16 sessions. Shaded region denotes frequencies from which elements were derived. **e.** Relative deviation in coherence length within elements vs. average coherence length across cortex. N = 4 mice, n = 16 sessions. Statistics determined by one-sample t-test comparing relative deviation of coherence length from 0. *** p<0.001; ns = not significant. AL: Anterolateral; AS: Active sensory; MS: Medial somatomotor; Visual: V; MA: Medial Associative

**Extended Data Figure 4:**
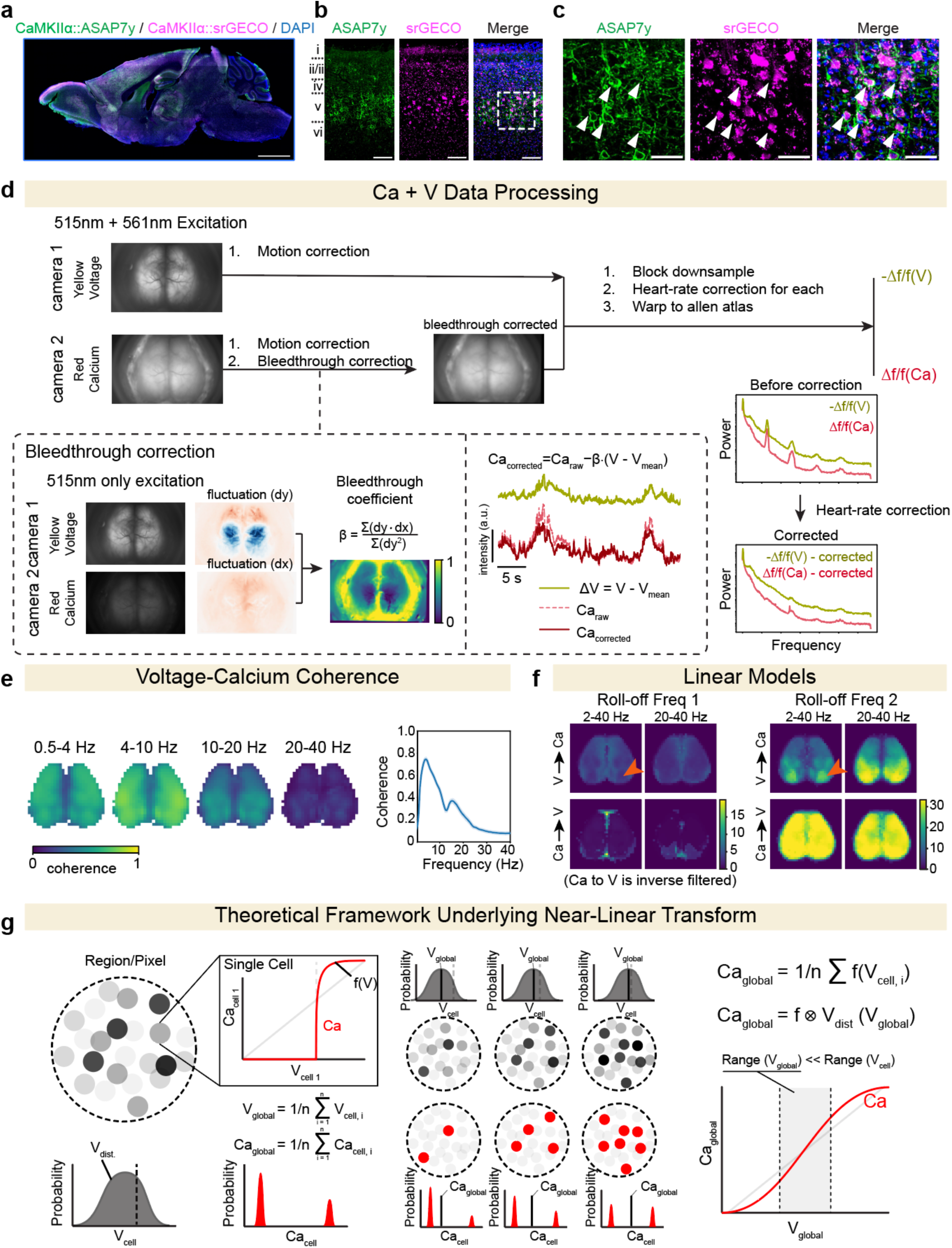
Measuring sub– and super-threshold dynamics and transformations. **a.** Histology of ASAP7y and srGECO co-expression. Representative sagittal section from a mouse following retro-orbital injection of PHP.eB-CaMKIIα::ASAP7y and PHP.eB-CaMKIIα::srGECO (scale bar: 2 mm). **b.** Cortical expression of ASAP7y and srGECO (scale bar: 100 μm). **c.** Zoomed-in view of layer V in b. Arrows denote cells co-expressing both indicators (scale bar: 50 μm). **d.** Processing simultaneous voltage (V) and calcium (Ca) data. Simultaneous V and Ca data is acquired with constant illumination at both 515nm (for ASAP7y) and 561nm (for srGECO), with emission split at 561 nm onto two cameras (camera 1: yellow/voltage channel; camera 2: red/calcium channel). Both channels undergo motion correction; the calcium channel additionally undergoes bleedthrough correction (dashed box). To correct for ASAP7y signal leakage into the red channel, a calibration session is recorded using only 515nm illumination at the start of each imaging session. Pixel-wise bleedthrough coefficients (β, spatial map shown in center) are computed as β = Σ(*dy* ⋅ *dx*)/Σ(*dy*^2^), where dy and dx are pixel-wise fluctuations in the yellow and red channels respectively. The corrected calcium signal is obtained as *Ca_corrected_* = *Ca_raw_* − β ⋅ (*V* − *V*^ˉ^), removing time-varying voltage artifacts while preserving the calcium baseline (example traces, center right). Both corrected channels then undergo block-downsampling, heart-rate correction, and warping to the Allen Brain Atlas (right). Power spectra before and after correction (bottom right) confirm removal of the heartrate harmonic from both the voltage and calcium channel. **e.** Map of the coherence between voltage and calcium across frequency bands. **f.** Parameters of linear models of Ca-V transforms in Fig 3d. Maps show the frequency of each of 2 poles (f1, f2) (or inverse poles in Ca➜V) for the 2-40Hz transforms and the 20-40Hz transforms. The V ➜ Ca (forward) is transform fit with two low pass filters, whose 3dB rolloff frequencies (poles) are plotted here. Arrows denote a sharp boundary around V1 where the optimal filters change, suggesting a potentially unique V ➜ Ca dynamics in V1. The Ca ➜ V (reverse) transform is fit with two inverse low pass filters (regularized to avoid blowing up), whose 3dB rolloff frequencies (zeros) are plotted here. **g.** Theoretical model explaining the linearity of the measured Ca-V transform– specifically, how non-linear single cell dynamics yield linear global signals. Each pixel averages the activity of many cells, each having a non-linear V➜Ca transform and a sharp spiking threshold. In a local circuit, neurons exhibit a distribution of membrane voltages. As the mean of this distribution shifts, the number of spiking cells changes continuously. Mathematically, the global V➜Ca transform is the convolution of the single-cell transform with the local voltage distribution. Because the global voltage range is much less than local voltage (∼20% Δf/f in an OEG pixel vs 100% Δf/f in single cell), the global transform in the explored range is approximately linear.

**Extended Data Figure 5:**
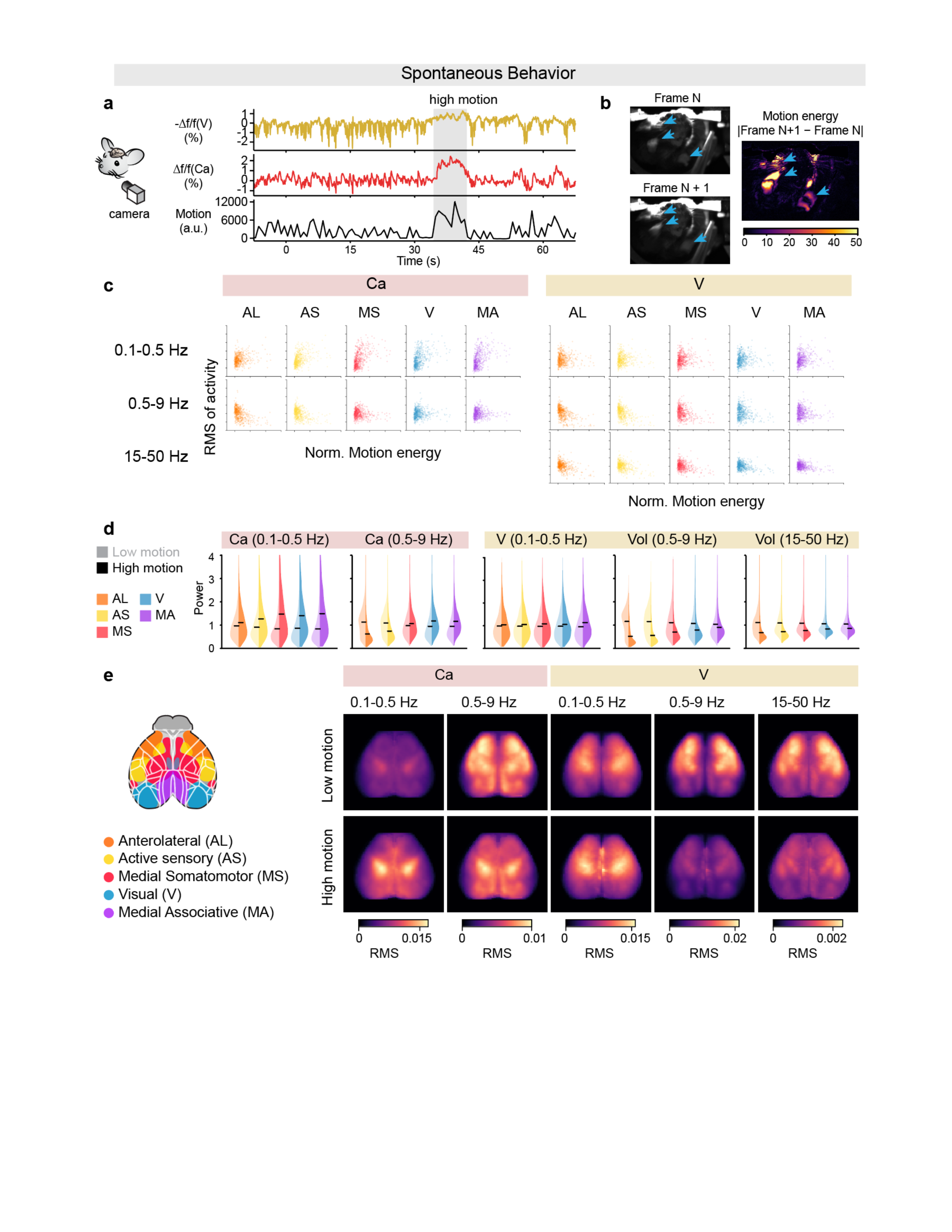
Modulation of cortical activity during spontaneous behavior. **a.** Representative traces of voltage (-Δf/f(V)), calcium (Δf/f(Ca)), and facial motion energy during spontaneous behavior, with a high-motion epoch highlighted (gray shading). **b.** Motion energy calculation. Motion energy is computed as absolute pixel-wise difference between consecutive frames from the behavior camera (100 Hz) (|Frame N+1 − Frame N|). Blue arrows highlight orofacial movement. **c.** Scatter plots of band-limited root-mean-square (RMS) activity within each functional motif versus normalized motion energy, computed in 1-s rolling windows, for calcium (left) and voltage (right) across three frequency bands. Low-frequency Ca and V activity (0.1-5Hz) is positively correlated with motion energy. Intermediate (0.5-9Hz) Ca show divergence between motifs while intermediate (0.5-9Hz) and high-frequency (15-50Hz) V is negatively correlated with motion energy. **d.** Violin plots comparing band-limited power within each functional motif during low-motion (bottom 75%, light) and high-motion (top 25%, dark) epochs. Both Ca and V show increased activity at low frequencies (0.1–0.5 Hz) during high motion across all motifs. At high frequencies (15–50 Hz), V activity decreases during high motion across all motifs. At intermediate frequencies (0.5–9 Hz), Ca shows increased activity in MS, V, and MA, whereas V shows decreased activity across all motifs. **e.** Spatial RMS maps across cortex for each frequency band and motion condition. At 0.1–0.5 Hz, high-motion epochs show broadly elevated RMS, particularly in the MS. At 0.5–9 Hz, Ca maps show decreased activity anteriorly (AL, AS) and increased activity posteriorly (MS, MA, V) during high motion, while V maps show broad decreases at both 0.5–9 Hz and 15–50 Hz. N =4 mice, n= 16 sessions. AL: Anterolateral; AS: Active sensory; MS: Medial somatomotor; Visual: V; MA: Medial Associative

**Extended Data Figure 6.**
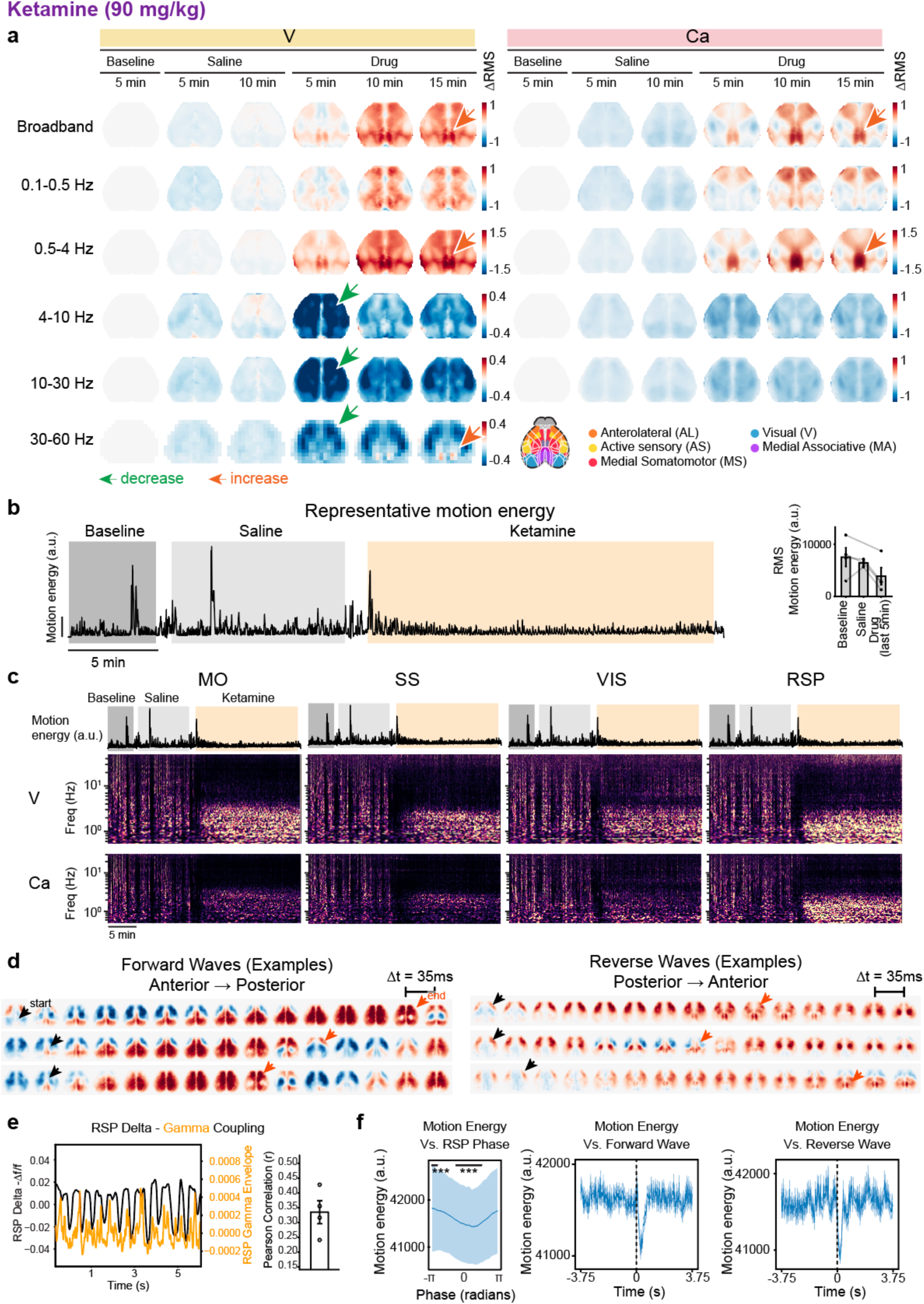
Ketamine-induced RSP-dominant oscillations and cortical waves. **a.** Band limited power in V and Ca across sessions. Global V activity at 90 mg/kg ketamine is increased in the 0.1–4 Hz band, with a particularly strong signal in RSP from 0.5-4Hz (orange arrow). This signal in V activity is accompanied by a 0.5-4Hz elevation in Ca activity most prominent in RSP, with a smaller signal in motor cortex at this dose. Higher frequency activity (> 4Hz) is heavily attenuated in both V and Ca with the exception of an increase in gamma (30-60Hz) power in RSP for V activity (beyond the temporal resolution of Ca measurement, orange arrow). Maps shown reflect mean data from N=4 mice. Green arrows highlight a significant suppression in the anterior cortex in 4-60 Hz voltage, and orange arrows highlight a significant increase in RSP in 0.5-4 Hz and 30-60 Hz. **b.** Left: Representative motion energy during baseline, saline, and ketamine. Right: Quantification of RMS motion energy across conditions (baseline, saline, and ketamine; measured during the last 5 minutes, 15 minutes post-injection). Ketamine trended toward reduced motion energy relative to baseline, though this did not reach statistical significance. N=4 mice. **c.** Example whitened spectrograms across cortex for ROIs in motor, somatosensory, visual, and RSP cortex; frequency is plotted on a log scale. Drug onset leads to an initial quieting across all bands followed by elevated power in the 0.1–4Hz band. Power above this band remains reduced except for gamma power especially in RSP. **d.** Example maps of forward (anterior to posterior) and reverse (posterior to anterior) waves; single trials identified by the protocol in Fig 4h. Black arrows mark the onset of the wave, and orange arrows mark the end of the wave. **e.** Coupling between delta-band and gamma-band power in RSP. Under ketamine, the RSP exhibits a delta-band 0.5-4 Hz oscillation (black) and a corresponding modulation in the 30-60 Hz (gamma) band (orange; amplitude of the Hilbert-transformed 30-60 Hz RSP signal). These two signals exhibited a Pearson correlation of 0.34, indicating that the gamma-band envelope is temporally correlated with the delta-band signal (N=4 mice). **f.** Coordination of oscillations/waves with mouse orofacial motion energy. *Left*: average and standard deviation of motion energy across RSP oscillation phase revealed that motion energy is phase-linked to the oscillation (1877 oscillation cycles, bar shows phases where the motion energy is significantly different than a scrambled control with p < 0.001 Bonferroni corrected). *Middle* and *right*: trial averaged motion energy during forward (132 waves) and reverse waves (76 waves), respectively, revealed motion reduction during the wave. (Representative mouse from N = 4 mice).

**Extended Data Figure 7:**
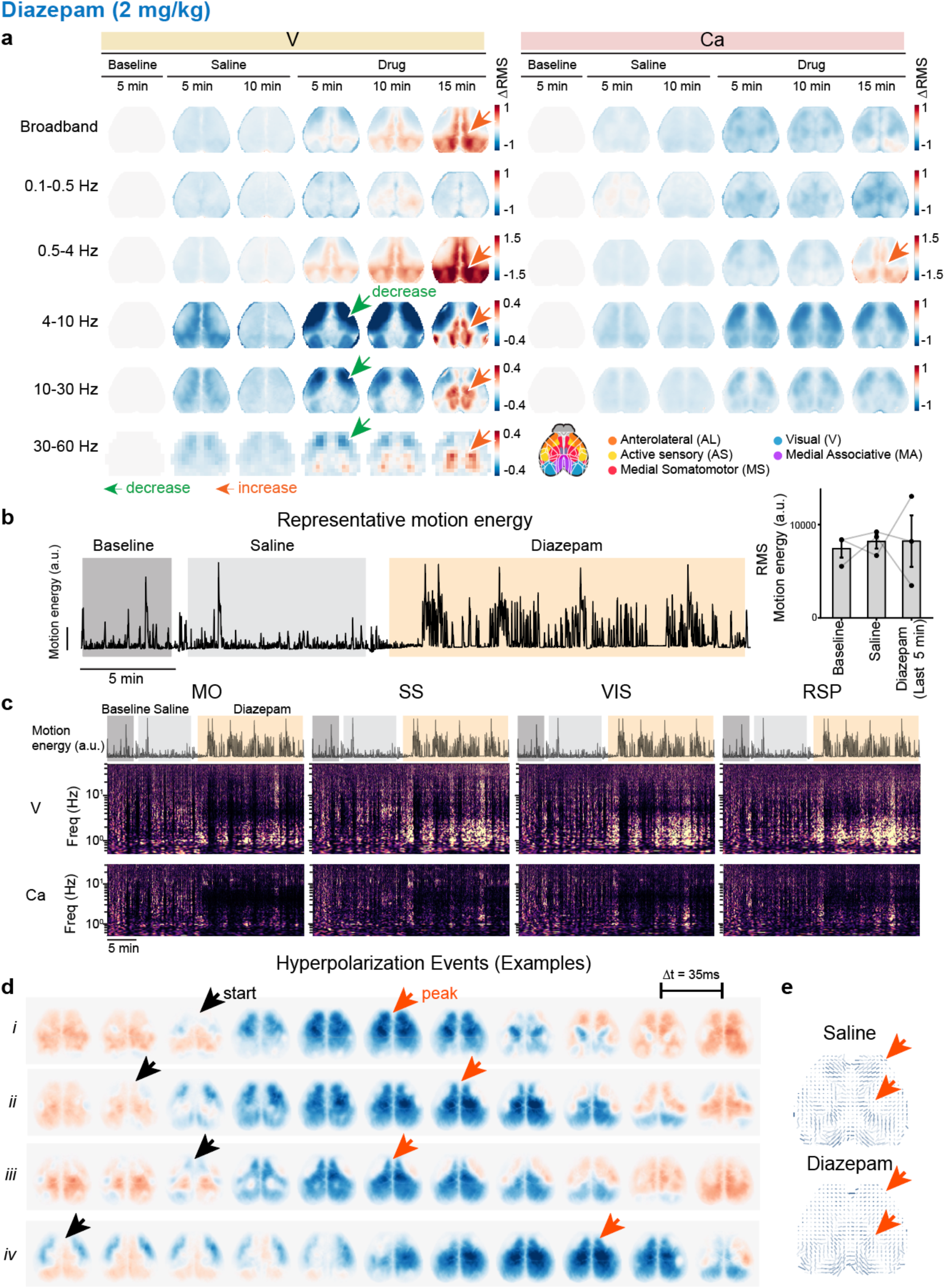
Diazepam induces cortex-wide activity modulation and hyperpolarization events. **a.** Band limited power in voltage (V) and calcium (Ca) signals across sessions. Plots show average maps across N = 3 mice. Both V and Ca maps showed increased activity from 0.5-4Hz (delta power) in visual and medial associative elements (orange arrows), with the effect more prominent in V. Outside of this band Ca activity was decreased globally, mirrored by V, although the V maps additionally revealed a diazepam-induced broadband high frequency elevation along the midline (orange arrows). V activity was attenuated in anterolateral and active sensory elements, and increased in the medial associative element at higher frequencies (green arrows)). **b.** Left: Representative motion energy during baseline, saline, and diazepam. Right: Quantification of RMS motion energy across conditions (baseline, saline, and diazepam; measured during the last 5 minutes, 15 minutes post-injection). N=3 mice. **c.** Representative whitened spectrograms across cortex in ROIs from motor, somatosensory, visual, and RSP cortex. Spectrograms are characterized by alternating periods of broadband high activity and quiescence, with broadband activation occurring preferentially during stationary epochs (low motion energy). **d.** Example single-trial examples of hyperpolarization events as in Figure 4g. Black arrows mark the start of the event, and orange arrows mark the peak of the event. Note that the four example events shown are of different propagation speeds. **e.** Example wave map during saline and diazepam. Noting the less directional waves in the anterior and midline areas (red arrows). Consistent for N =3 mice.

**Extended Data Figure 8:**
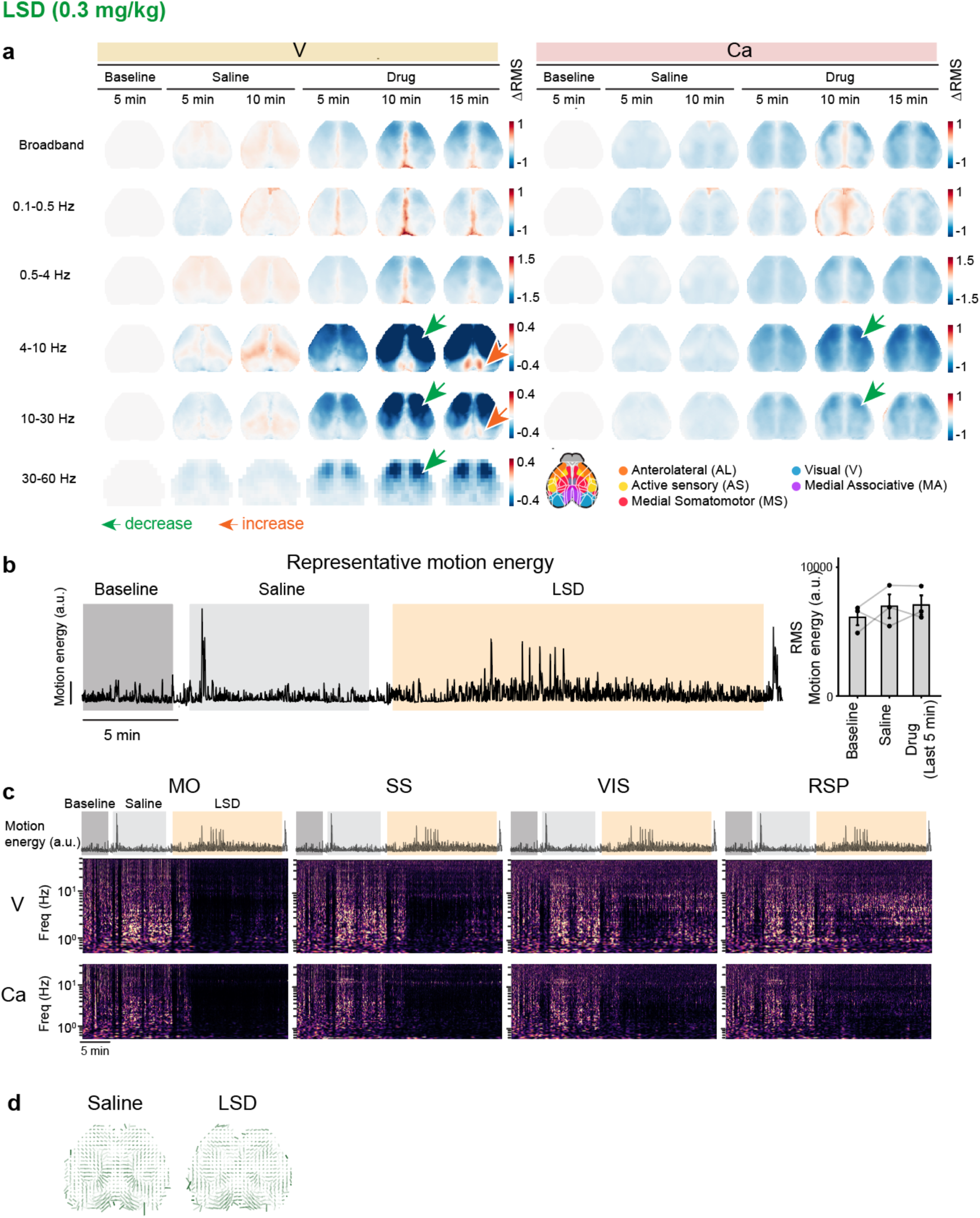
LSD induces cortex-wide activity modulation. **a.** Band limited power in voltage (V) and calcium (Ca) across sessions. Plots show average maps across N = 3 mice. All frontal elements were powerfully suppressed in LSD, especially at higher frequencies (>4 Hz) (green arrows). RSP showed a band limited increase in power from 4-10Hz, detectable only in V (orange arrows). An infraslow midline elevation visible in both V and Ca likely reflects hemodynamic contributions given its spatial correspondence with cortical vasculature. **b.** Left: Representative motion energy during baseline, saline, and LSD. Right: Quantification of RMS motion energy across conditions (baseline, saline, and LSD; measured during the last 5 minutes, 15 minutes post-injection). N=3 mice. **c.** Example whitened spectrograms across cortex from ROIs in motor, somatosensory, visual, and RSP; frequency plotted on a log scale. Drug onset led to immediate broadband quieting. Intermediate frequency (4-10Hz) activity was relatively less affected in visual cortex and RSP. **d.** Example wave map during saline and LSD. Consistent for N =3 mice. No significant differences between conditions.

**Extended Data Figure 9:**
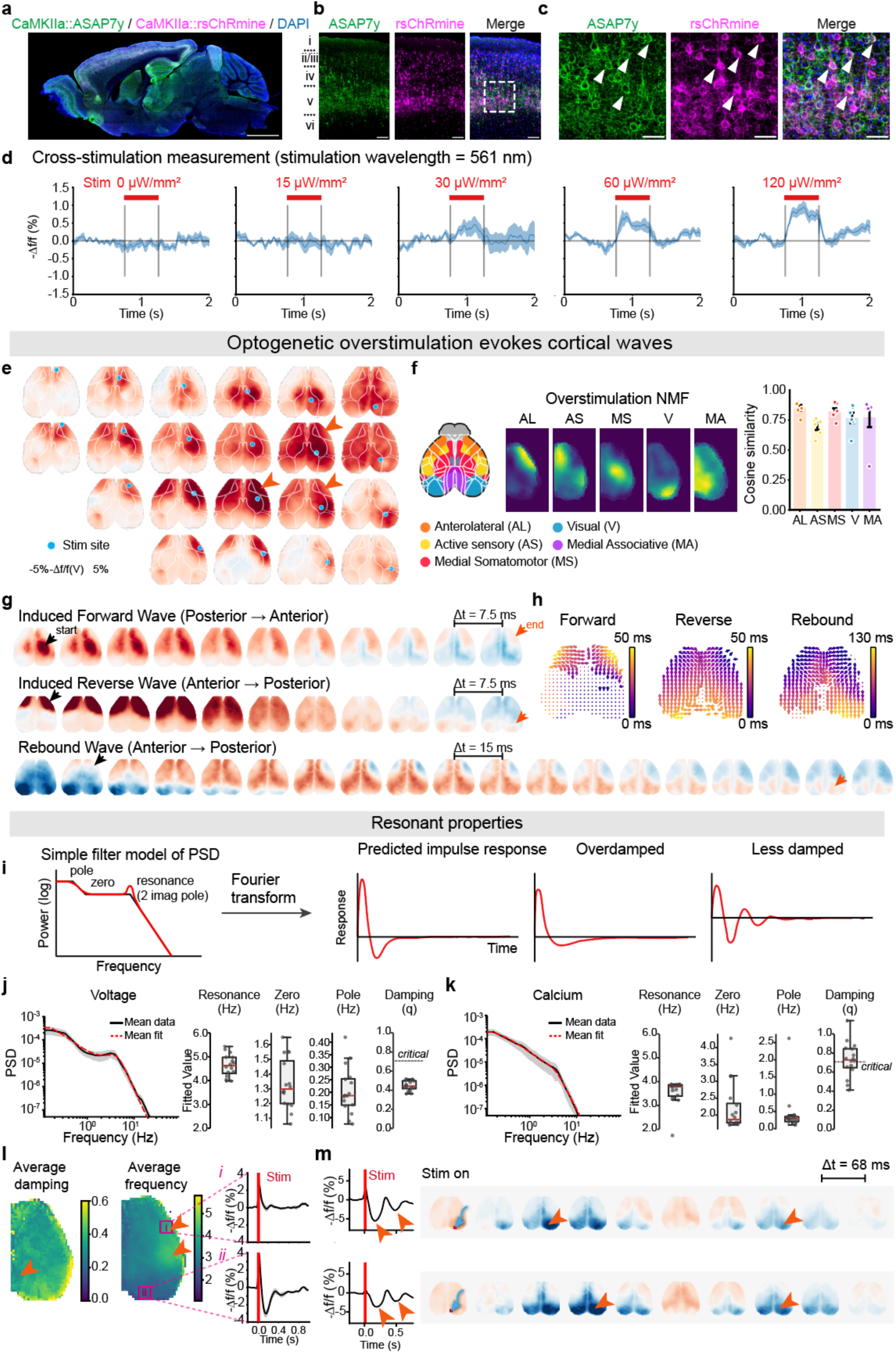
Causal tests of cortical organizational principles. **a.** Histology of ASAP7y and rsChRmine co-expression: representative sagittal section following retro-orbital injection of PHP.eB CaMKIIα::ASAP7y and PHP.eB CaMKIIα::rsChRmine-oScarlet (scale bar: 2 mm). **b.** Cortical expression of ASAP7y and rsChRmine-oScarlet (scale bar: 100 μm). **c.** Zoomed-in view of layer V. Arrows denote cells co-expressing both constructs (scale bar: 50 μm). **d.** Cross-stimulation measurement. Trial averaged time traces of cross-stimulation measurement from Fig 5c. During this measurement, a 3mm circle of 561nm light (homogenized with a multimode fiber) was projected unilaterally onto the somatosensory cortex continuously for 500ms. Plots show the response measured as the trial averaged – Δf/f averaged across mice. Filled area: s.e.m. across N = 5 mice. **e.** Overstimulation (via calculated 20 mW/mm^2^ optogenetic-stimulus light power density at target) breaks through network barriers to induce global waves. Representative evoked responses (45 ms post-pulse) shown, demonstrating recruitment of large, cross-element cortical areas (element boundaries overlaid in white). Arrows denote AL element recruitment from distal stimulation sites. **f.** Non-negative matrix factorization (NMF) of overstimulation responses decomposes cortical activity into the same elements identified during spontaneous activity that functionally parcellate cortical activity. *Bottom*: Cosine similarity between evoked recruitment patterns and the established functional parcellation; N = 6 mice. Although overstimulation recruits multiple areas, this suggests that the recruitment still happens in an element-wise fashion, with multiple elements being recruited by one stimulus. **g.** Overstimulation triggers forward (from anterior to posterior as most commonly seen with ketamine) and reverse (from posterior to anterior) global cortical waves. A secondary “rebound” wave following the reverse wave propagates at reduced velocity. Example wave shown; all waves visible in supplemental video 5. **h.** Wave propagation and velocity analysis of optogenetically induced global waves. Flow fields show consistent propagation structure that proceeds across the cortex monotonically in time. **i.** Model used to fit power spectra in Figure 5l; each spectrum is fit with a pole, zero, and a resonance (frequency and damping factor). Shown are impulse responses predicted from this model from the average fit (q = 0.5), overdamped fit (q = 0.9), and less damped fit (q = 0.2). **j.** Calculated resonance frequency and damping from overstimulation points positioned in an unbiased screen spanning cortex. Arrows denote a slightly reduced average damping in RSP, and higher resonance frequencies in barrel and anterolateral cortices. *Right*, measured average impulse responses at 2 example points (i, ii); average of N = 6 mice. **k.** Overstimulus-induced local evoked responses (in RSP and V1) exhibit prolonged, damped oscillations. Arrows denote hyperpolarization maxima. *Right:* Whole cortex maps of damped oscillations revealed oscillatory activity in both visual and medial associative networks. Blue arrows denote stimulation sites, orange arrows denote hyperpolarization maxima. Representative from N = 5 mice).

**Extended Data Figure 10:**
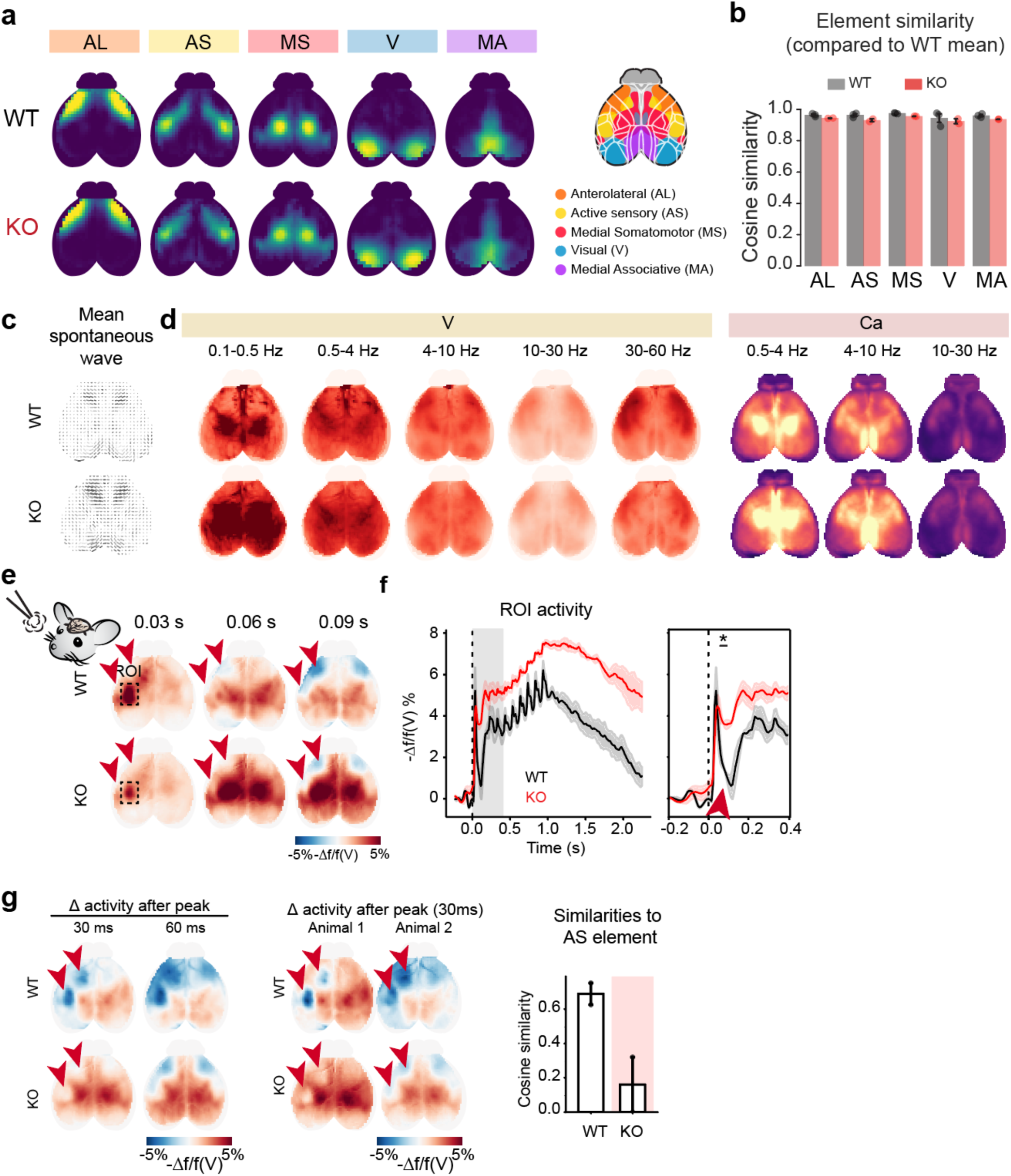
CIM voltage mapping reveals a high speed inhibitory deficit in the Cntnap2 model of autism. **a.** Functional assessment of spontaneous and evoked cortical dynamics in *Cntnap2* knockout (KO) and wild-type (WT) littermates expressing ASAP7y via retro-orbital injection of AAV-PHP.eB CaMKIIα::ASAP7y. **b.** Elemental structure appeared conserved in spontaneous cortical dynamics, as networks derived via NMF of all-to-all pixel-wise coherence (1–10 Hz) revealed no significant spatial reorganization between WT and KO as quantified by cosine similarity, statistics determined by two-tailed t test. **c.** Stability of local wave propagation. Analysis of local lag vectors and propagation anisotropy revealed no major restructuring of spontaneous wave trajectories in the *Cntnap2* KO vs. control. **d.** Comparative power distribution. RMS power across frequency bands was found to be consistent between WT and KO cohorts in both voltage (V) and calcium (Ca) imaging (Ca data from a separate cohort expressing AAV-PHP.eB-CaMKIIα::GCaMP8m; N=6 WT (n=59 12-min sessions), N=6 *Cntnap2* KO (n=60 12-min sessions). **e.** Testing sensory-evoked changes: trial-averaged responses to whisker air puff stimulation (10 Hz). In WT mice, sensory stimulation induced rapid activity followed by suppression within the active sensory (AS) network (arrows). This suppressive response was nearly abolished in *Cntnap2* KO mice (N = 2 mice per group), revealing a deficit in fast-timescale cortical inhibition. **f.** Temporal profile of activity within the AS network (dashed ROI in e). *Left*: Mean fluorescence traces from the indicated ROI showed sustained elevation (lack of desensitization) in KO mice compared to WT. *Right*: Zoomed view of the initial response phase. In WT mice, a sharp inhibitory dip (red arrowhead) followed the initial evoked peak, which was greatly reduced in KO mice (N = 2 per group), two-tailed Welch’s t test, pointing to a specific deficit in fast time-scale cortical inhibition. **g.** Spatial localization of inhibitory deficits. *Left*: Maps of activity change “Δ activity”) 30 and 60 ms after the peak response. In WT mice, suppression specifically recruited the ipsilateral and contralateral anterior sensory (AS) elements (red arrowheads). This recruitment was significantly reduced in KO mice. *Middle:* Maps of activity change 30 ms after peak response in all mice. *Right*: Cosine similarity between the puff-evoked suppression pattern (30 ms post peak) and the established AS element. *Cntnap2* KO mice exhibited significantly lower similarity.

**Supplemental Video 1: CIM recording of ASAP7y**.

Example recording of CaMKIIα::ASAP7y across the entire dorsal cortex. Colorbar is scaled from –5% to 5% –Δf/f, and video is played at 1/3x speed (111 FPS movie of 333 FPS recording).

**Supplemental Video 2: Toy models of cortical dynamics**.

Example videos of the toy models of cortical dynamics. i) Stochastic local oscillator model. ii) Unconstrained propagation model. iii) Compartmentalized propagation model.

**Supplemental Video 3: Simultaneous voltage and calcium recording**.

Example recording of simultaneous CaMKIIα::ASAP7y (voltage) and CaMKIIα::srGECO (calcium) across the entire dorsal cortex. Colorbar is scaled from –5% to 5% –Δf/f for ASAP7y and –3% to 3% Δf/f for srGECO, and video is played at 1x speed (133 FPS movie of 133 FPS recording).

**Supplemental Video 4: Sub– and super-threshold recording under drug-induced states**.

Example recording of pharmacologically induced cortical dynamics. Each column shows simultaneous CaMKIIα::ASAP7y (voltage) and CaMKIIα::srGECO (calcium) across the entire dorsal cortex of the same mouse under saline, ketamine, diazepam, and LSD. Colorbar is scaled from –5% to 5% –Δf/f for ASAP7y and –3% to 3% Δf/f for srGECO, and video is played at 1x speed (133 FPS movie of 133 FPS recording).

**Supplemental Video 5: An unbiased all-optical electrophysiology screen**.

Trial-averaged recordings of single pulse optogenetic stimulation screens from a representative mouse CaMKIIα::ASAP7y (voltage indicator) and CaMKIIα::rsChRmine (opsin). Blue dots show stimulation sites and timing. Frames are interpolated during these stimulation frames. Weak stimulation is plotted from –3% to 3% –Δf/f and strong stimulation from –5% to 5% –Δf/f. Video is played at 1/10x speed (13 FPS movie of a 133FPS recording).

## Methods

### Experimental Model and Subject Details

All procedures were performed in accordance with protocols approved by the Stanford University Institutional Animal Care and Use Committee (IACUC) and the National Institutes of Health guidelines. Experiments utilized male and female C57BL/6J mice (Jackson Laboratory, 000664) and *Cntnap2* knockout (*Cntnap2*^−/−^ or *Cntnap2^KO^*) and wild-type (*Cntnap2^+/+^* or *Cntnap2^WT^*) mice (Jackson Laboratory, 017482). *Cntnap2^KO^* and *Cntnap2^WT^*mice were generated from in-house heterozygous-by-heterozygous breeding to provide littermate controls. Animals were aged 8–12 weeks at the time of surgery and group-housed under a standard 12-hour light/dark cycle. Post-surgery, subjects were transitioned to a 12-hour reversed light cycle for all subsequent experiments.

### Viral Vectors and Transduction

All viral vectors were packaged at the Stanford University Viral Core. The following constructs were utilized: PHP.eB CaMKIIα::ASAP7y-Kv2.1 (1×10^11^ vg/mouse), PHP.eB CaMKIIα::srGECO (5×10^11^ vg/mouse), and PHP.eB CaMKIIα::rsChRmine-oScarlet-Kv2.1 (1×10^11^ vg/mouse), PHP.eB CaMKIIa::GCaMP8m (7×10^11^ vg/mouse). Systemic viral delivery was performed via retro-orbital injection in mice aged 4–5 weeks.

### Surgery

Procedures were performed under sterile conditions. Mice were anesthetized with 4-5% isoflurane and maintained with 1–2% isoflurane and administered sustained-release buprenorphine (0.5mg/kg) subcutaneously for analgesia. Following scalp removal and skull cleaning, a custom headplate was affixed using dental cement. The skull was sealed with a thin layer of cyanoacrylate glue and clear nail polish. Mice recovered for at least one week prior to experimentation. For electrophysiological recordings, craniotomies (AP +1.6 mm, ML 1.0 mm) were performed one day prior to recording under isoflurane anesthesia. Exposed sites were cleaned with saline and protected with Kwik-Cast (WPI). A reference electrode was positioned over the olfactory bulb and secured with dental cement.

### Conformal immersion microscopy

#### System overview and excitation path

A widefield macroscope was constructed for high-speed, dual-channel cortical imaging. The system employed two synchronized sCMOS cameras (Teledyne Photometrics) synchronized via Streampix software (NorPix) with acquisition rates >500 Hz. Excitation light was provided by 510 nm and 560 nm LEDs (Thorlabs) filtered via 10 nm bandpass filters and combined through a 550 nm dichroic mirror. The primary optical path used an 85 mm f/1.2 objective (Nikon). Data acquisition rates were tailored to the specific experimental requirements: spontaneous activity in mice expressing ASAP7y alone was acquired at 333 or 400 Hz, while dual-channel imaging (ASAP7y and srGECO) and recordings in autism-model mice were performed at 133 Hz.

#### Conformal immersion and field-flattening optics

To maximize photon collection, the 85 mm objective (Nikon, ZY Optics) was coupled to a 0.58x focal reducer (speed booster, Metabones), yielding an effective aperture of f/0.7. To compensate for the restricted depth of field and cortical curvature, a custom dual-axis aspheric field-flattener lens was fabricated from optical-grade resin via stereolithography and polished. Glycerol (n = 1.47) served as the immersion medium between the lens and the cranial window.

#### Spectral separation

The emission path utilized a quad-band dichroic mirror (Chroma ZT440/514/561/640rpc) and a quad-band emission filter (Chroma ZET440/514/561/640m). A subsequent 560 nm dichroic mirror split the signal into yellow (lambda < 561 nm) and red (lambda > 561 nm) channels for simultaneous detection.

### Optogenetic Stimulation

Optogenetic stimulation was delivered using a 561 nm laser (Coherent, OBIS) directed by a 2D micro-electromechanical system (MEMS) mirror. The MEMS assembly was integrated into the excitation path via a 3 mm collimating lens and a removable mirror. Laser positioning was controlled via custom Python software. High-intensity stimulation used 15 ms pulses at 20 mW (at focal plane); low-intensity stimulation used 4 mW. Timestamps were synchronized to imaging data via a Nidaq board (National Instruments) and Streampix. This modular design enabled spatially patterned laser stimulation of the cortex without compromising the light-collection efficiency of the wide-field fluorescence path.

### Sensory stimulation

Tactile or visual stimuli were delivered unilaterally via an Arduino-controlled solenoid valve (airpuff to whisker pad) or blue LED (470 nm). Stimulus trains (10 Hz) were controlled by an Arduino and synchronized to image acquisition via TTL pulses recorded in StreamPix.

### Behavioral Recording

Mice were habituated to head-fixation for at least three days prior to data collection. During experimental sessions, animals were allowed to settle for 5 min after head-fixation before recordings. Spontaneous behavior was recorded in darkness using a high-speed infrared camera capturing facial motion, synchronized to the sCMOS imaging clock via Streampix software (NorPix). During optogenetic stimulation sessions, low-level ambient light was provided to saturate retinal activity and minimize visually evoked potentials in the cortex.

### Electrophysiological Recordings

Local field potential (LFP) was acquired using Neuropixels 1.0 probes and SpikeGLX software (AP gain = 500, using the bottom 384 electrode sites). Before insertion, probes were cleaned with Tergazyme (Alconox), rinsed with deionized water, and coated with CM-DiI (Thermo Fisher). Probes were inserted acutely at a 45° to a depth of 4mm. Comparison LFP was taken approximately 300um from the cortical surface. A 10-min stabilization period followed insertion before data acquisition began. Probe synchronization was acquired via a NIdaq board (National Instruments) and aligned using TPrime. Imaging data were synchronized by sending TTL triggers per frame to the same NIdaq.

### Drug administration

Pharmacological agents included ketamine (MWI/VetOne, NDC: 13985-584-10, ketamine HCl, 90 mg/kg), LSD (RTI International, (+)-lysergic acid diethylamide (+)-tartrate (2:1), 0.3 mg/kg), and diazepam (Natco Pharma USA, NDC: 69339-137-05, 2 mg/kg). For all imaging sessions, recordings consisted of a 5-min baseline, followed by 10 min post-saline injection and 20 min post-drug injection. Saline and drugs were administered subcutaneously during active recording. To account for injection-induced motion artifacts, frames spanning 0.5 s before to 60 s after injection were excluded from subsequent analysis.

### Histology and immunohistochemistry

Mice were deeply anesthetized and transcardially perfused with 4% paraformaldehyde (PFA) in PBS. Brains were post-fixed in 4% PFA overnight at 4°C and subsequently sectioned sagittally at a thickness of 50 um using a vibratome. For immunolabeling, free-floating sections were incubated with a rabbit anti-GFP conjugated to Alexa Fluor 488 (Invitrogen; 1:750 dilution) in blocking buffer overnight at 4°C. Sections were counterstained with DAPI and mounted. Representative images were acquired using a confocal microscope to verify fluorophore expression and anatomical localization.

### Data Preprocessing

#### Motion correction and dF/F calculation

Imaging data were motion corrected using the Enhanced Correlation Coefficient algorithm (cv2.MOTION_EUCLIDEAN)^77^. Relative fluorescence was calculated as ΔF/F = (F – F0) / F0, where F0 represents the mean fluorescence of each pixel over the temporal dimension. Following these steps, the resulting ΔF/F data were spatially downsampled by a factor of 4.

#### Spectral crosstalk correction

Following motion correction and timestamp alignment using Streampix metadata, spectral crosstalk was corrected post-acquisition. Yellow-to-red leakage was quantified for each subject during a reference period under 510 nm excitation only. A pixel-wise bleedthrough coefficient, *k*, was calculated via least-squares: *Red_raw_*(*t*) = *k* ⋅ *Yellow_raw_*(*t*). Red channel data were subsequently corrected by linear subtraction: *Red_corrected_*_’_(*t*) = *Red_raw_*(*t*) − *k* ⋅ *Yellow_raw_*(*t*)

#### Cardiac Artifact Removal

Cardiac artifacts were removed using pixel-wise regression similar to previous publication^78^. A reference signal was extracted from cortical regions identified by high signal variance and bandpass filtered (10–14 Hz) to isolate the cardiac rhythm. Instantaneous phase was extracted via Hilbert transform and smoothed. A matrix of four harmonics was utilized to regress out cardiac components from individual channel.

#### Atlas registration

Processed images were warped to the Allen Mouse Brain Common Coordinate Framework (CCFv3)^38^ for standardized spatial analysis. Registration was performed using an affine transformation to align the experimental field of view with the dorsal cortical projection of the atlas^79,34^. Pixels falling outside the defined anatomical boundaries of the Allen framework were excluded from subsequent analysis.

#### Behavioral Analysis

Oral-facial movement was quantified as motion energy (temporal derivative of behavioral ROIs containing the mouse face), 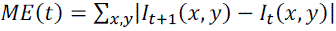, and synchronized to imaging timestamps via linear interpolation.

### Local field potential extraction

LFPs were extracted from the wideband Neuropixels data from the top of the probe using the ecephys_spike_sorting pipeline^80,81^. Raw data were processed through the catGT helper module to concatenate trials, apply high-pass filtering to the AP band, and isolate the LFP band via low-pass filtering at 300–500 Hz. LFP data were synchronized to behavioral and imaging timestamps using the tPrime_helper module to correct for clock drift between the Neuropixels acquisition system and external NIDAQ-recorded triggers. Reduced coherence at high frequencies between LFP and optical signals likely arise from the cell-type specificity of the voltage indicator. While the indicator signal depends directly on membrane voltage, the LFP contains contributions from both membrane potentials and extracellular dipole effects^26^.

### Computational methods

Spectral analyses were restricted to frequencies below the Nyquist limit (fps/2) for each dataset.

#### Band-limited RMS analysis

To characterize the spatial distribution of cortical activity across different frequency regimes, dF/F signals were first bandpass filtered into discrete frequency bands. For each subject, the RMS amplitude was calculated pixel-wise for each band as the square root of the temporal mean of the squared signal.

#### Pixel Frequency Profiling

Spectral energy distribution was quantified pixel-wise using a continuous wavelet transform (CWT) with variable-cycle Morlet wavelets. The wavelet cycle count was linearly scaled from 3 at 0.1 Hz to 10 at 150 Hz. Time-frequency power (*S*) was calculated as the squared magnitude of the coefficients and normalized by the temporal mean to yield *S_norm_*. To assess state-dependent spectral shifts, frames were partitioned into high-activity (top 25th percentile) and low-activity (bottom 75th percentile) states based on the sum of *S_norm_* across all frequencies. A “pixel frequency profile” was then generated by calculating the fraction of the total spectral energy in each state at each frequency. These profiles were computed for all pixels within the Allen CCFv3 mask and averaged across the field of view.

#### Resonance characterization

Spontaneous power spectra were fit using a transfer function combining a first-order filter with a single-mode resonator to capture both the broadband 1/f slope and the specific resonant peak:

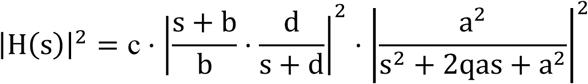

where *s* = *i*2π*f*. This allowed for the simultaneous estimation of the natural frequency (*a*), damping ratio (*q*), and the pole/zero frequencies that characterize the low frequency spectrum.

For optogenetics evoked oscillation, the post-stimulus signal decay in ROIs surrounding the stimulation points (a 4×4-pixel region centered on each stimulation coordinate) was modeled as a damped harmonic oscillator:

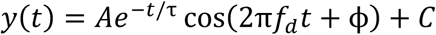

where *f*_d_ is the damped frequency and τ is the decay time constant. Natural frequency (*a*) and damping ratio (*q*), were derived using the following relations:

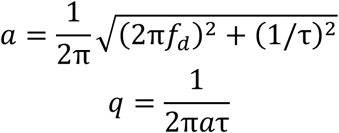

#### Traveling wave analysis

##### Wavefront velocity and directionality

Local propagation dynamics were quantified by computing velocity vectors (v_x_, v_y_) from dF/F data bandpass filtered at 5–50 Hz. Signals were spatially downsampled (2×2) and temporally upsampled by a factor of 9 to achieve sub-frame lag resolution. Velocity was derived from the peak cross-correlation lag between each pixel and its four cardinal neighbors, computed within a sliding window of 1350 samples (post-upsampling) with 50% overlap. Local trajectories were summarized as covariance ellipses, where eccentricity and orientation represent directional anisotropy and preferred propagation axes across the cortex. We measure ellipticity as the ratio 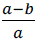 where a is the covariance of the first principle component and b is the covariance of the second. We measure the Bhattacharyya coefficient between distributions assuming normal distributions and using the definition:

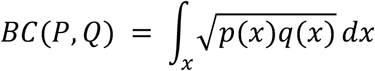

where P and Q are two distributions with probability density p and q respectively and x integrates the 2D plane. To calculate this coefficient we use the closed form for multivariate normal distributions.

##### Ketamine induced wave propagation and phase-event coupling

Propagation patterns during ketamine-induced waves were characterized by sampling signal intensities along equidistant linear trajectories (oriented at 50° from the horizontal) across the right half of the imaging field (excluding the retrosplenial cortex, RSP). Spatiotemporal profiles were generated by averaging signal intensities across each trajectory over time. Discrete wave events were defined as sequential 0-to-1 transitions across linear regions of interest (ROIs). The relationship between wave timing and the global RSP oscillatory state was determined by extracting the instantaneous phase—calculated via the Hilbert transform—at the onset and offset of detected propagation events. Phase distributions were visualized as polar histograms to assess the stability of wave initiation relative to the underlying rhythm.

#### Calcium Voltage linear transform

Transforms were estimated by fitting 2^nd^ order linear filters to the data using minimization over parameters with L-BFGS. The V ➜ Ca (forward) is transform fit with two low pass filters, and scale parameter:

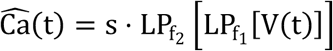

Where s is the scale factor and f1 and f2 are the 3dB roll-off frequencies. The Ca ➜ V (reverse) transform is fit with two inverse low pass filters regularized to avoid blowing up:

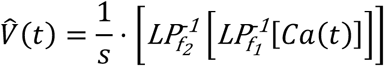

Where s is the scale factor and f1 and f2 are the 3dB roll-off frequencies.

#### Coherent length analysis

##### Distance-dependent coherence

Spatial scales of functional connectivity were quantified by estimating spectral coherence (C_xy_) as a function of Euclidean distance (*d*). To reduce computational load, Monte Carlo sampling with utilized with increasing resolution until convergence. For each dataset, pixel pairs (∼500 pairs) were randomly sampled at separations ranging from 1 to 50 pixels using a circular shell search algorithm (0.5-pixel tolerance). Coherence was calculated via Welch’s method^82^

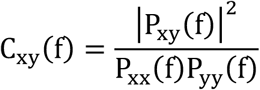

##### Coherent length estimation

The characteristic spatial correlation length λ(*f*) was defined as the furthest distance at which the normalized coherence (normalized to the value at the minimum sampled distance) remained above 1/*e*. This value was estimated for each frequency bin using linear interpolation across the distance-coherence matrices.

#### Coherence-based non-negative matrix factorization (NMF) decomposition

To identify spatially distinct functional networks that leads to the decreased coherence length, we performed NMF on the spectral coherence features of the dF/F signal. Data were spatially binned (2 × 2) and pixel-wise coherence was calculated relative to the global mean using Welch’s method as above. Coherence stacks were then partitioned into frequency bands: 0.1–4 Hz, and the resulting coherence matrix V was decomposed into components such that *V* ≈ *WH*, where *W* represents the spatial membership maps and *H* the component signatures. To ensure a reasonable number of factors, we increased factor count until elements began splitting bilaterally (n = 6). Each pixel was assigned a discrete label corresponding to the component with the highest membership value, yielding topographic maps of frequency-specific activity elements.

